# Substrate Profiling of RNF216 Uncovers a Translation-Linked OTUD4 Regulatory Axis

**DOI:** 10.64898/2026.08.26.747332

**Authors:** Wei Wei, Ruochuan Liu, Jing Zhang, Shu Liu, Antoinette J. Charles, Devansh G. Asati, Zachary D. Allen, Delance Wright, Kangli Peng, Ellissa Krekeler, Nima Mosammaparast, Jun Yin, Angela M. Mabb

**Author notes:** Corresponding Authors: Angela M. Mabb, Georgia State University, 100 Piedmont Ave. SE, Atlanta, GA 30303, USA, Jun Yin, Georgia State University, 50 Decatur St. SE, Atlanta, GA 30303, USA.

## Abstract

Mutations in the E3 Ubiquitin (Ub) ligase *RNF216* cause Gordon Holmes syndrome (GHS), a neurodegenerative disorder accompanied by neuroendocrine disruption. We developed an orthogonal ubiquitin transfer (OUT) platform to capture RNF216 substrates in neuronal cells and identified OTUD4, a deubiquitinating enzyme (DUB) mutated in GHS, and FMRP, a neuronal-enriched translational repressor. RNF216 predominantly synthesizes K6-linked Ub chains on OTUD4 to induce its degradation, forming donut-shaped structures in neurons. In return, OTUD4 removes the ubiquitination of RNF216 and FMRP. Analysis of RNF216 substrates revealed biological functions regulating protein synthesis, a shared function of the OTUD4-RNF216 substrate interaction network. Indeed, RNF216 expression increased protein synthesis rates while *Rnf216* deletion decreased dendritic development in neurons. Overall, our findings show that RNF216 and OTUD4 balance rates of protein synthesis and degradation and suggest GHS-related mutations in *RNF216* or *OTUD4* may offset this balance, triggering neurodegeneration.

**Highlights:**

- Orthogonal Ubiquitin Transfer cascade identifies RNF216 substrates in neuronal cells
- The deubiquitinating enzyme OTUD4 is a RNF216 substrate
- RNF216 controls rates of protein synthesis and degradation
- RNF216 and OTUD4 operate as a catalytic pair

## Introduction

Protein ubiquitination, the covalent addition of a 76-residue ubiquitin (Ub) to cellular proteins, is an essential posttranslational modification (PTM) that regulates a variety of biological processes^1–3^. Ub transfer to cellular targets is carried out by an enzymatic cascade consisting of an Ub-activating enzyme (E1), Ub-conjugating enzyme (E2), and Ub ligase (E3). The human genome encodes an abundant (> 600) and diverse class of E3 Ub ligases^4,5^ that drive substrate selectivity in cells. Importantly, numerous mutations in the E3 ligase machinery have been identified in various pathological conditions, especially in neurological disorders^6^. Ubiquitination is reversed by deubiquitinating enzymes (DUBs), which remove Ub conjugates from E3 substrate proteins^7^. However, there is a large gap in understanding the variety of substrates targeted by E3 Ub ligases and DUBs in the nervous system.

Although multiple neurological diseases are associated with mutations in Ub-transferring and processing enzymes^6^, there is one type of neurological disorder, Gordon Holmes syndrome (GHS), that is almost exclusively caused by mutations in E3s and DUBs regulating protein ubiquitination^8^. GHS is an autosomal recessive adult-onset neurodegenerative disorder that is characterized by reproductive dysfunction, ataxia, dementia, and neurodegeneration^8^. Mutations in the Ub machinery that include E3s *RNF216* and *CHIP/STUB1* alongside the DUB *OTUD4* are linked to GHS^8–14^. Loss-of-function mutations in *RNF216,* including mutations that disrupt its E3 catalytic activity, have been discovered^8,15–17^. Digenic mutations within the E3 catalytic RBR domain of RNF216 and outside of the DUB domain of OTUD4 (G333V equivalent to G398V in the large isoform) have also been identified^8^. *OTUD4* mutations occur in congenital hypogonadotropic hypogonadism, which has some phenotypic overlap with GHS^18^. RNF216 levels are also reduced in Alzheimer’s disease, where a primary point of pathology is reduced protein degradation^19^.

In this study, we developed an orthogonal ubiquitin transfer (OUT) cascade to enable the exclusive transfer of an engineered UB (xUB) through RNF216 to its substrates for their specific capture and identification by proteomics. The OUT screen assembled an RNF216 substrate profile of 174 targets, which included FMRP, a neuronal translational repressor^20–22^ and the DUB OTUD4^23^. Here, we find that RNF216 predominantly assembles K6-linked Ub chains on OTUD4, which induces the formation of donut-like structures in neurons and decreases OTUD4 stability. On the other hand, OTUD4 counteracts the E3 Ub ligase activity of RNF216 to remove Ub chains from RNF216 and FMRP. The OTUD4 GHS mutant (G398V) has tempered DUB activity, and deletion of *Otud4* in mice mirrors GHS phenotypes previously found in *Rnf216* knockout (KO) mice^24,25^. Modulation of RNF216 alters global protein synthesis rates and decreases the complexity of neuronal dendrites, which rely heavily on protein synthesis for their development^26^.

Taken together, our findings suggest that the E3-DUB partnering between RNF216 and OTUD4 fine-tune the ubiquitination of substrate ensembles, where RNF216 functionally operates as a molecular “proteostat” to regulate protein degradation and synthesis in neuronal cells. Our work also highlights that GHS is a disorder resulting from not only disrupted protein degradation but also of imbalanced protein synthesis.

## Results

### Engineering the OUT cascade to identify RNF216 substrates

We first sought out to better understand the range of RNF216 substrates. The OUT cascade enables the exclusive transfer of an engineered Ub (xUB) with R42E and R72E mutations through an engineered xE1-xE2-xE3 cascade to the substrates of a specific E3 in the cell (Figure 1A)^27,28^. The OUT method assigns E3 substrates by tracing the direct transfer of xUB from the E3 to its substrate proteins. This is advantageous relative to indirect readouts that are hampered by changes in protein stability or ubiquitination levels that could be influenced by factors such as nonproteolytic Ub chain conjugation to substrates and E3s affecting proteasome activity or the expression and Ub ligase activity of other E3s. Modeling off the Parkin OUT system^29^, we engineered the RBR domain of RNF216 to enable its transfer of xUB for profiling its substrates in neuronal cells (Figures S1A-S1B).

**Figure 1:**
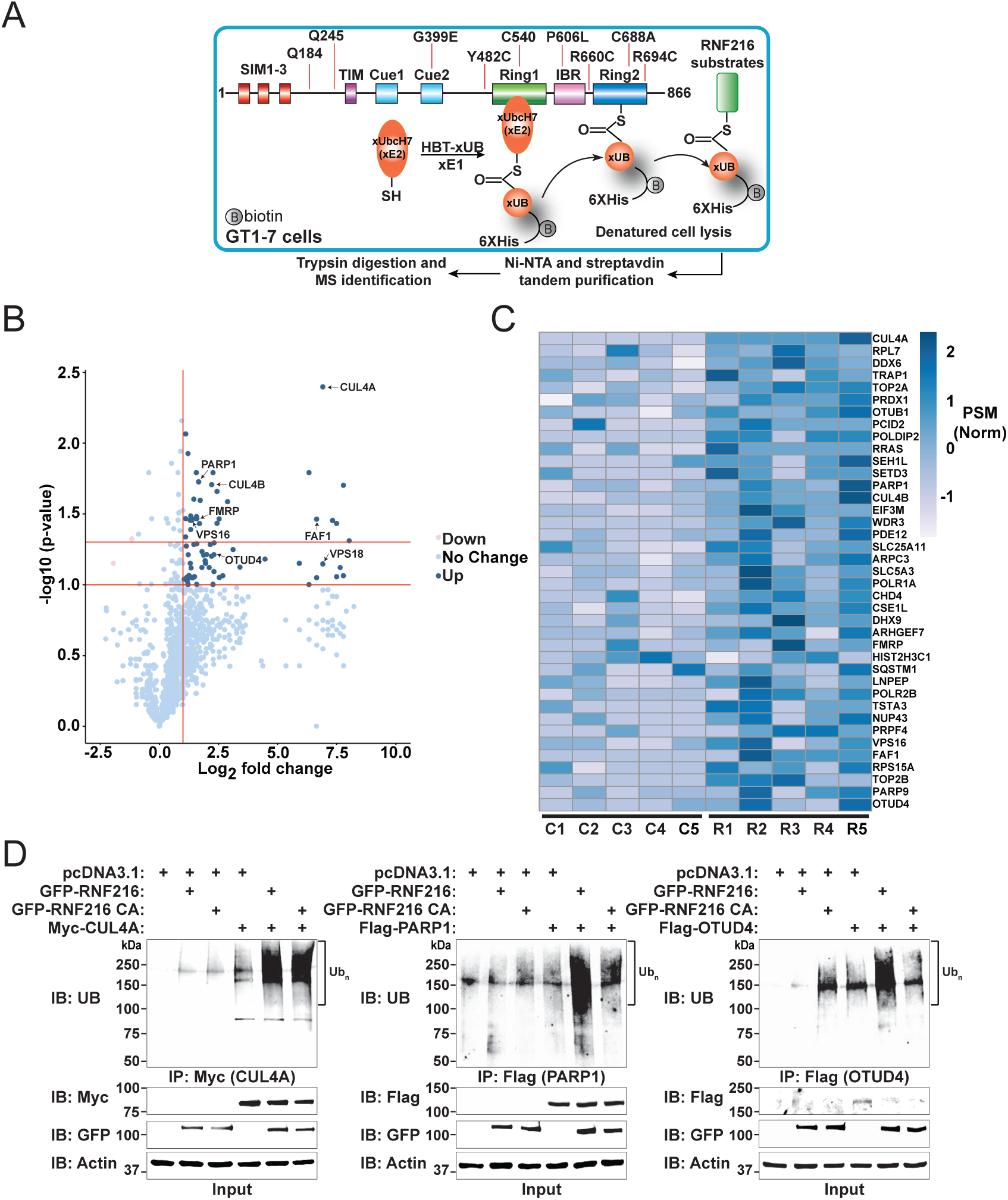
An engineered Orthogonal Ubiquitin Transfer (OUT) cascade identifies RNF216 substrates in mouse hypothalamic cells. (A) Schematic of OUT cascade for RNF216. xUB is transferred through the orthogonal engineered xE1-xE2-xE3 (xRNF216) cascade to substrates of xRNF216 in GT1-7 cells. xUB-conjugated proteins are tandem purified with the 6×His-biotin tag (HBT) fused to xUB (HBT-xUB) under denaturing conditions to identify the direct substrates of xRNF216. (B) Volcano plot of RNF216 substrates identified using RNF216 OUT. The dark blue dots show the differential expression of targets in xRNF216 compared to catalytic inactive xRNF216 CA, Log 2 [PSM ratio xRNF216/ xRNF216 CA] >1 and -Log 10 p > 1. N = 5 independent biological replicates. (C) Heatmap for significant targets as calculated using normalized PSM values from OUT. 5 replicates from the control cells (C1-C5) expressing xRNF216 CA; 5 replicates (R1-R5) from cells expressing xRNF216. (D) Validation of top hits from OUT by co-expressing Myc-CUL4A (p = 0.004), Flag-PARP1 (p = 0.02), and Flag-OTUD4 (p = 0.06) with WT RNF216 or RNF216 CA mutant. HEK 293 cells were transfected with respective plasmids. Cell lysates were subjected to denaturing immune purification with anti-Myc or anti-FLAG antibodies and processed samples were immunoblotted with an anti-UB antibody. Inputs were immunoblotted with anti-Myc, -GFP, -Flag or -Actin antibodies. N = 3 independent biological replicates.

The RBR domain at the C-terminus of RNF216 consists of RING1, in-between RING (IBR) and RING2 subdomains, with RING1 engaging the E2∼Ub thioester conjugate and RING2 harboring the catalytic Cys (C688) that bridges Ub transfer to the Lys residues on substrates. The engineered E1-E2 pair (xUBA1-xUBCH7) is incompatible with wild-type (WT) RNF216 (Figure S1C). The crystal structure of the RNF216 RBR in complex with the UbcH7-Ub conjugate captured the active conformation of the RBR domain during Ub transfer from UbcH7 to RBR, in which the C-terminus of Ub runs in parallel to a β-strand in RING2 to approach C688, the catalytic Cys of the RBR of RNF216^30^. Residues I683, S685, and E686 in the β-strand of RING2 are near R42E and R72E mutations in xUB and may block xUB from reacting with C688 of the RBR domain (Figure S1B). We constructed several RBR mutants of RNF216 by replacing I683, S685, or E686 individually, or their combinations with Arg to match the R42E and R72E mutations in xUB. We found that I683R or the double mutants of I683R/S685R and I683R/E686R reacted with xUB transferred through the xUBA1-xUBCH7 pair, with the I683R/E686R mutant showing the highest reactivity in xUB loading (Figure S1D). We thus incorporated the I683R/E686R double mutation into full-length RNF216 and used it to assemble an xUBA1-xUBCH7-xRNF216 OUT cascade for transferring xUB to RNF216 substrates in the cell.

To express the OUT cascade of RNF216 in neuronal cells, we constructed a 2-plasmid system, with the cDNAs of xUBA1 and xRNF216 cloned in the pCAGGS vector and cDNAs of xUBCH7 and HBT-xUB cloned in a pAAV vector containing a P2A bicistronic element at the juncture of the two cDNAs for their expression as individual proteins in cells (Figure S7A)^31^. These two plasmids were cotransfected into mouse hypothalamic immortalized cells (GT1-7), which were selected due to our previous findings that these cell types are affected in our *Rnf216* KO mouse model (Figure S7A)^24^. We validated the expression of Flag-xUBA1, V5-xUBCH7, myc-xRNF216, and HBT-xUB in GT1-7 cells, and purified HBT-xUB-conjugated proteins from cells expressing the RNF216 OUT cascade for proteomic identification (Figure S7A). In addition, we included a C688A mutation (CA) in the xRNF216 gene in the pCAGGS vector and used it as a negative control (control cells). The purification of HBT-xUB conjugated proteins from these two groups of cells were performed five times using tandem Ni-NTA and streptavidin affinity columns, and proteins bound to the streptavidin resin were digested by trypsin for identification by LC-MS (Figure S7B). Proteins with 2-fold or higher peptide intensities purified from OUT cells compared to the control cells were assigned as potential substrates of RNF216 (Figure 1B). Proteomics analysis demonstrated a total of 174 targets at low stringency cutoff (P < 0.1) and 54 targets at high stringency cutoff (P < 0.05) (Figures 1B, 1C). Targets in order of p value (high to low stringency) included CUL4A, PARP1, and OTUD4, which were validated using WT RNF216 and its catalytic inactive (CA) mutant for Ub assays in HEK293 cells, demonstrating the high fidelity of the OUT method to identify RNF216 substrates (Figure 1D).

### OTUD4 is regulated by RNF216-mediated ubiquitination

From the RNF216 substrate list, OTUD4 was a notable validated target that stood out from our screen (Figures 1B and 1C), given that an *OTUD4* mutation (G398V) alongside an *RNF216* mutation was identified in GHS (Figure S2A) and *Otud4* manipulations also exhibit a neuronal phenotype in zebrafish knockout models^8^. In mouse brain, OTUD4 expression was highest during early development with a decrease beginning around puberty, whereas RNF216 expression was relatively consistent (Figure S2B). Given these relationships, we sought to further validate OTUD4 as an RNF216 substrate, which included elucidating a potential mechanism of regulation. OTUD4 was ubiquitinated by RNF216 in vitro using recombinant RNF216 and immunopurified OTUD4 from HEK293 cells (Figure 2A). Using Tandem Ubiquitin Binding Entities (TUBEs) to capture polyubiquitinated proteins, we found that OTUD4 ubiquitination was reduced in *Rnf216* GT1-7 KO cells (Figure 2B)^24^. The reduction in OTUD4 ubiquitination was coupled to an increase in its stability as OTUD4 levels remained elevated in GT1-7 *Rnf216* KO cells, as shown in an anisomycin pulse-chase assay (Figure 2C). There was a close-to-significant increase in the steady-state level of OTUD4 in *Rnf216* KO GT1-7 cells (Figure 2D). *Rnf216* KO brains had decreased OTUD4 ubiquitination and elevated steady-state levels of OTUD4 in the hypothalamus of *Rnf216* KO mice (Figure 2E and 2F).

**Figure 2:**
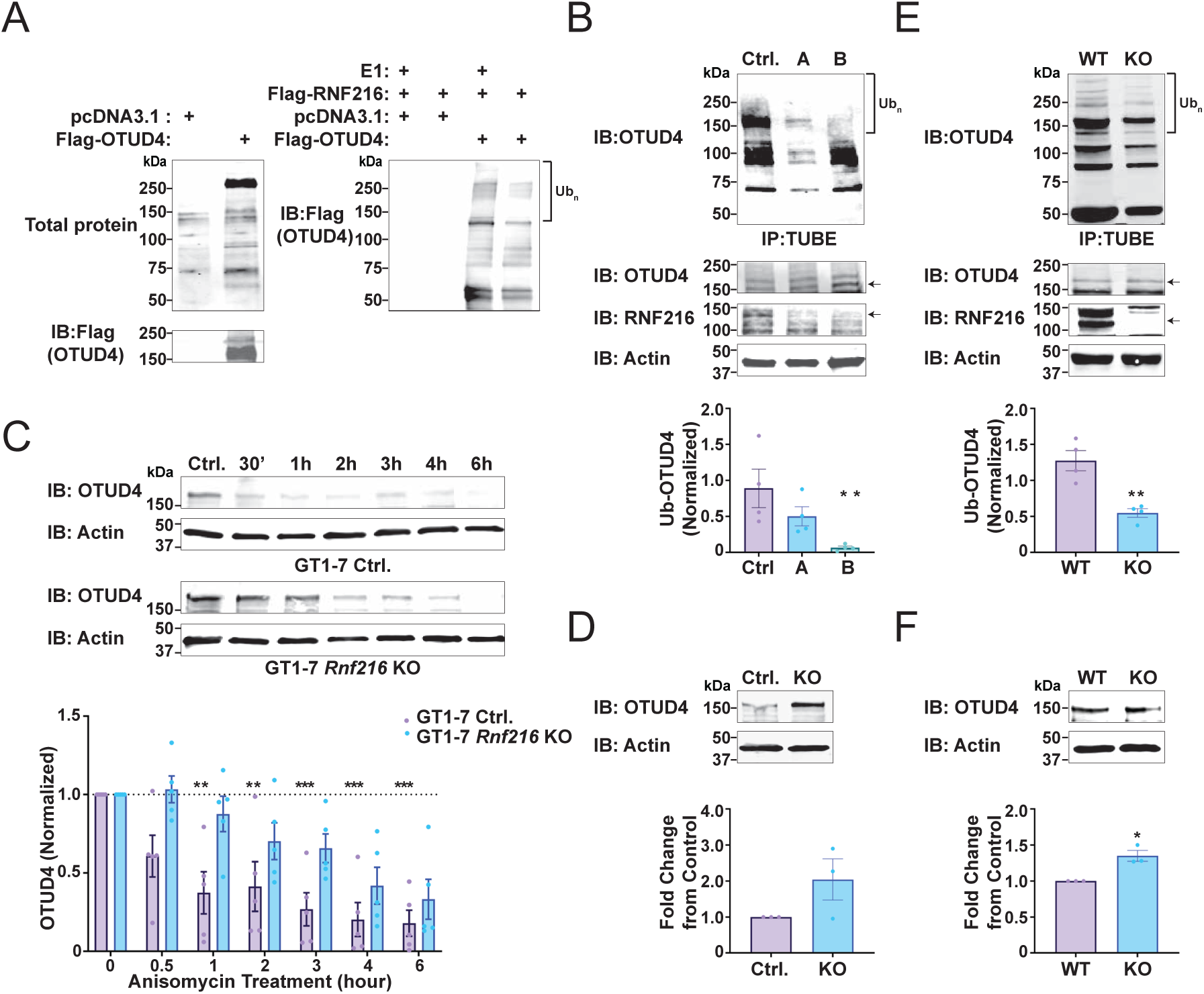
RNF216 is required for OTUD4 ubiquitination and is destabilizing. (A) In vitro ubiquitination of OTUD4 by RNF216. *Left*, Flag-OTUD4 was expressed in HEK 293 cells and immunopurified using native, stringent conditions. *Right*, Immunopurified Flag-OTUD4 was subjected to an in-vitro ubiquitination reaction using recombinant Uba1 (E1), UbcH7 (E2), Ubiquitin (Ub), and full-length Flag-RNF216. Reactions were probed using an anti-Ubiquitin (Ub) antibody. (B) *Top*, GT1-7 cells with control (Ctrl.), partial (A), and full loss of RNF216 (B) were measured for changes in OTUD4 ubiquitination using Tandem Binding Ubiquitin Entities (TUBEs). Processed samples were immunoblotted with an anti-OTUD4 antibody. Inputs were immunoblotted with anti-OTUD4, -RNF216, or -Actin antibodies. *Bottom*, Quantification of OTUD4-UB conjugates. * p = 0.02, One-way ANOVA post hoc Tukey’s t-test. N = 4 independent biological replicates. (C) Extended half-life of OTUD4 in *Rnf216* KO GT1-7 cells. Crispr control and *Rnf216* KO (KO) GT1-7 cells were plated and incubated with vehicle (DMSO) or anisomycin (20 µM) for 30’, 1h, 2h, 3h, 4h, or 6h. Samples were immunoblotted with anti-OTUD4 and -Actin antibodies. *Bottom*, Quantification of OTUD4 in Ctrl. and KO GT1-7 cells. ** p < 0.05, *** p < 0.005. One-way ANOVA with Dunnett’s multiple comparisons test. N = 5 independent biological replicates. (D) Near significant elevation of OTUD4 in *Rnf216* KO cells. Welch’s t test, P = 0.056. N = 3 independent biological replicates. (E) OTUD4 ubiquitination is reduced in mouse *Rnf216* KO adult brain. *Rnf216* WT and KO whole brain tissue lysates were measured for changes in OTUD4 ubiquitination using TUBEs. *Bottom*, Quantification of OTUD4-UB conjugates. **p = 0.003, unpaired t-test. N = 4 independent biological replicates. (F) Modest increase in OTUD4 in *Rnf216* KO brain. Quantification of OTUD4 large isoform. N = 3, Welch’s t test, * p = 0.043. N = 3 independent biological replicates.

To further gain insight into how the GHS (G398V) mutation in OTUD4 might alter its ubiquitination by RNF216, we performed assays in HEK293 cells using combinations of Ub configurations. As previously shown, RNF216 was able to enhance the ubiquitination of OTUD4 in this system (Figure 3A). However, this ubiquitination was reduced in the OTUD4 GHS mutant. Mutation of the OTU catalytic site of OTUD4 (C45A), which is critical for its DUB activity^32^, increased RNF216-mediated ubiquitination compared to WT (Figure 3A). These data demonstrate the opposing effects of the OTUD4 GHS mutation G398V and the catalytically inactivating mutation C45A on RNF216-mediated OTUD4 ubiquitination.

**Figure 3:**
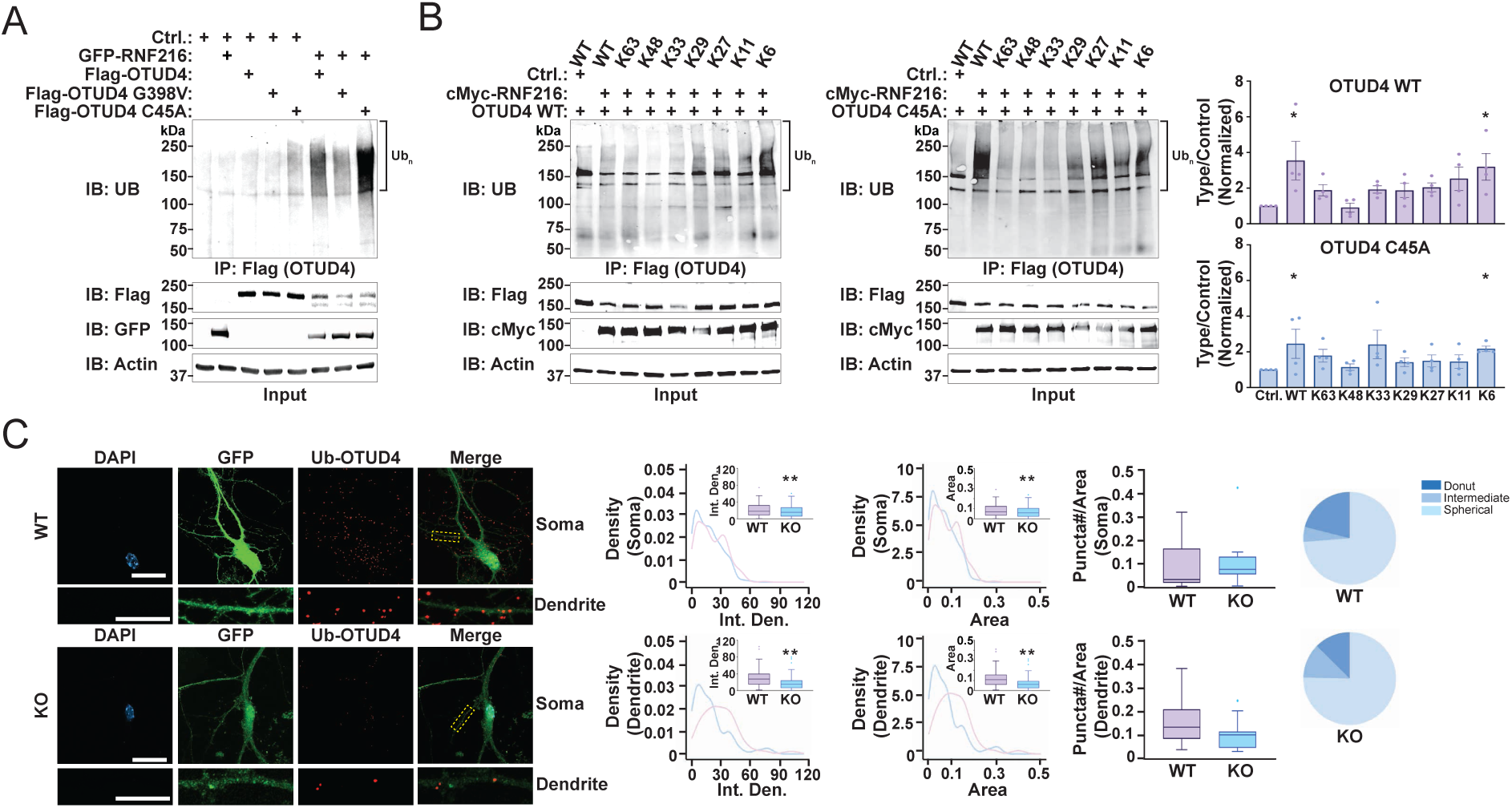
RNF216 dominantly forms K6-Ub linkages on OTUD4 and forms Ubiquitin-OTUD4 donut-like structures in hippocampal neurons. (A) OTUD4 GHS mutant (G398V) has reduced RNF216-mediated ubiquitination whereas the OTUD4 catalytic inactive mutant (C45A) has enhanced RNF216-mediated ubiquitination. HEK 293 cells were transfected with Flag-OTUD4, the Flag-OTUD4 GHS-associated mutation (G398V) or catalytic inactive Flag-OTUD4 (C45A) in combination with GFP-RNF216. Cell lysates were immunopurified under denaturing conditions using an anti-FLAG antibody and processed samples were immunoblotted with an anti-UB antibody. Inputs were immunoblotted with anti-Flag, -GFP, or -Actin antibodies. (B) RNF216 dominantly assembles K6-Ub linkages on OTUD4. *Left*, HEK 293 cells were transfected with HA-UB and single Lysine Ub mutants with Myc-RNF216 and Flag-OTUD4 or Flag-OTUD4 C45A. Flag-OTUD4 was immunopurified under denaturing conditions and processed samples were immunoblotted with an anti-UB antibody. Inputs were immunoblotted with anti-Flag, -Myc, or -Actin antibodies. *Right*, Ub chain quantification for OTUD4 variants. One-way ANOVA. Dunnett’s test. P < 0.05. N = 4 independent biological replicates. (C) Deletion of *Rnf216* decreases Ub-OTUD4 complexes. *Rnf216* WT and *Rnf216* KO primary hippocampal neurons were transfected with GFP to outline neuron morphology. Neurons were fixed and then subject to PLA assay as stated in C. *Middle*, *Rnf216* KO neurons had reduced Ub-OTUD4 intensities. Comparison of puncta intensities and area per cell between *Rnf216* WT and KO was performed as in C. *Inset*, Box plots show comparison of mean values by unpaired t-test. *Right*, Pie chart showing a reduction in the formation of donut-like structures in *Rnf216* KO neurons. N = 21-22 cells per condition. Scale bars = 20 and 5 μm.

Previous studies suggest RNF216 catalyzes the formation of K11- and K63-linked Ub chains^17,19,33^. However, we found that K48-and K63-specific TUBEs were unable to capture ubiquitinated OTUD4 in HEK 293 cells (Figure S3A-C). These findings were not related to poor efficacy of TUBEs as the K63-specific TUBE was highly reactive towards RNF216, matching the dominant Ub chain linkage on self-ubiquitinated RNF216 (Figure S1A)^17^. These findings then compelled us to investigate RNF216-catalyzed OTUD4 ubiquitination using Ub mutants at various linkage sites. The single Lys mutants of Ub were coexpressed with RNF216 and OTUD4 in HEK293 cells, followed by immunopurification of OTUD4 to measure its ubiquitination level. Among all the single Lys mutants of Ub, the K6-only mutant generated the highest level of Ub chains on WT OTUD4 as well as the C45A mutant with masked DUB activity (Figure 3B, left and right). These findings demonstrate that RNF216 dominantly forms K6-linked Ub chains on OTUD4 with the possibility of forming other chain types, in line with the finding that RNF216 can assemble heterotypic ubiquitin chains on certain substrates^19^.

To gain insight into RNF216 and OTUD4 interactions, we used immunocytochemistry to resolve their spatial interactions in neurons. OTUD4 and RNF216 were found to colocalize in neuronal regions that included dendrites, axons, and the soma but not in the nucleus (Figure S4A). OTUD4 also colocalized with Ub signals but this localization was not significantly different in *Rnf216* KO neurons (Figure S4B). To obtain a more detailed interaction of OTUD4 with Ub, we used a proximity ligation assay (PLA) in combination with super resolution microscopy to trap and amplify OTUD4-Ub conjugates at the single molecule level. Interestingly, we observed 100 - 300 nm sized donut-like structures in primary hippocampal neurons (Figure 3C, right). Interestingly, deletion of *Rnf216* led to a reduction in the intensity, size, and donut-shape of Ub-OTUD4 structures in KO cells (Figure 3C). Altogether, our data support a role for RNF216 in regulating OTUD4 ubiquitination and complex assembly in neurons.

### Deubiquitination of RNF216 by OTUD4 is independent of RNF216 stability

Since RNF216 recognizes OTUD4 as a ubiquitination substrate, we assayed if OTUD4 catalyzes the deubiquitination of RNF216 in return as there is precedent that DUBs and E3 Ub ligases can cross-regulate one another under different contexts^34–37^. To consider this, we conducted in vitro DUB assays against RNF216 assembled Ub chains. The purified DUB domain of OTUD4 (1-180) was found to be catalytically active in cleaving K48-linked Ub chains (Figure S5A). We set up a self-ubiquitination reaction of RNF216 in vitro and found that the DUB domain of OTUD4 could remove Ub chains assembled on RNF216 after quenching the E3 ligase activity of RNF216 with apyrase or EDTA (Figure S5B). We also compared the activity of the WT DUB of OTUD4 with the catalytically inactive mutant (C45A) and found that the WT DUB of OTUD4 but not C45A could remove Ub chains on RNF216 (Figure 4A). Together, these findings show that OTUD4 can reverse RNF216 self-ubiquitination in vitro. To determine if OTUD4 could modulate RNF216 Ub chain assembly in cells, we expressed RNF216 with full-length OTUD4, the GHS mutant (G398V), or the catalytic inactive (C45A) mutant in HEK293 cells to examine their effects on RNF216 ubiquitination. Expression of OTUD4, but not the C45A mutant, reduced RNF216 ubiquitination. The OTUD4 G398V mutant caused an intermediate decrease in RNF216 self-ubiquitination, suggesting that its DUB activity may be slightly compromised or that interaction between OTUD4 and RNF216 is reduced (Figure 4B).

**Figure 4:**
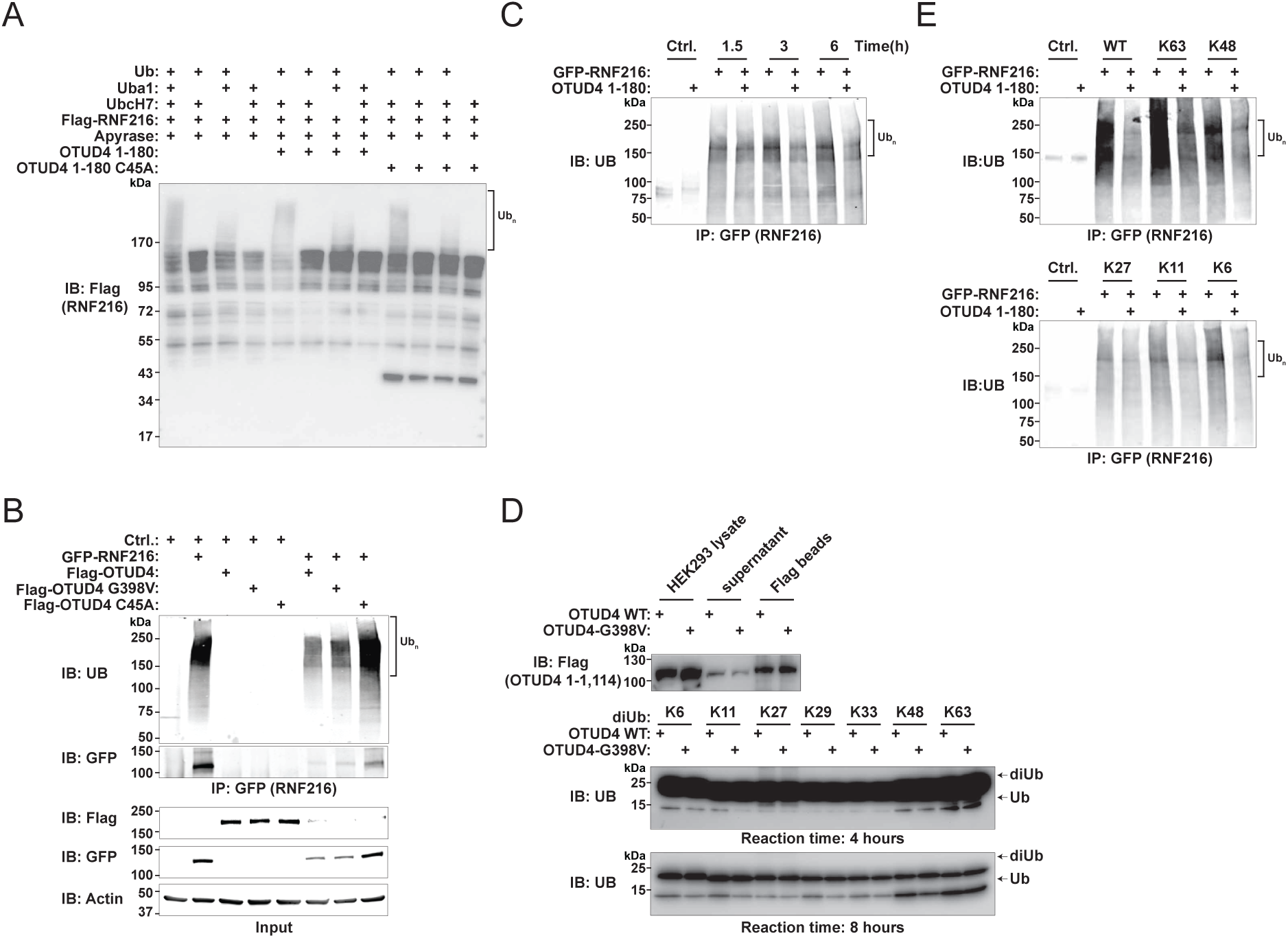
OTUD4 deubiquitinates RNF216 with a prevalence for RNF216 K63-Ub linkages in cells. (A) Recombinant DUB domain of OTUD4 disassembles RNF216 ubiquitinated chains in vitro. In vitro ubiquitination reaction using recombinant Ub, E1 (Uba1), E2 (UbcH7), Flag-RNF216 and recombinant DUB domain of OTUD4 WT or catalytic inactive (C45A) fragment (1-180). Samples were incubated as described in methods and reaction products were subjected to SDS-PAGE followed by immunoblotting with an anti-Flag antibody. (B) OTUD4 WT and the OTUD4 GHS-associated mutation (G398V) differentially deubiquitinate RNF216. HEK 293 cells were transfected with Flag-OTUD4, the Flag-OTUD4 G398V, and catalytic inactive Flag-OTUD4 C45A with or without GFP-RNF216. Cell lysates were immune purified under denaturing conditions using an anti-GFP antibody and processed samples were immunoblotted with an anti-UB antibody. Inputs were immunoblotted with anti-Flag, -GFP, or -Actin antibodies. (C) Recombinant OTUD4 disassembles RNF216 ubiquitinated chains. HEK 293 cells were transfected with GFP-RNF216 WT. Cell lysates were immunopurified under native conditions using an anti-GFP antibody, incubated with recombinant OTUD4 (1-180), and then washed as stated in the methods. Terminated reactions were immunoblotted with an anti-UB antibody. N = 3 independent biological replicates. (D) Full-length OTUD4 and the G398V mutant with a Flag tag were expressed and purified from HEK293 cells by immunoprecipitation with anti-Flag beads and elution from the beads with 3×Flag peptide. The activities of the WT OTUD4 and the G398V mutant were assayed by their cleavage of diUb of various linkages in reactions of four and eight hours. WT OTUD4 showed a higher activity in cleaving diUb linkages of K6, K11, and K48 than the G398V mutant, while the WT and G398V mutant OTUD4 have comparable activities in cleaving K63-linked diUb. (E) Recombinant OTUD4 (1-180) disassembles a variety of RNF216 self-assembled chain types with a dominant preference for K63-linkages. HEK 293 cells were transfected with GFP-RNF216 WT and HA-Ub or HA-Ub mutants containing only one Lysine. Cell lysates were immunopurified under native conditions using an anti-GFP antibody, incubated with recombinant OTUD4 (1-180), and then washed as stated in the methods. Terminated reactions were immunoblotted with an anti-UB antibody.

The DUB domain of OTUD4 was also active against immunopurified full-length ubiquitinated RNF216 (Figure 4C, Figure S5C). To determine the cleavage specificity of OTUD4 on Ub chains, we screened the activity of full-length OTUD4 in cleaving di-Ub conjugates of various linkages and found WT OTUD4 immunopurified from HEK293 cells mainly cleaves K48 and K63-linked Ub chains with a preference of cleaving K63-linked Ub chains over the K48-linked chains (Figure 4D). This specificity of OTUD4 for Ub chain linkages matches a previous report showing that phosphorylation of OTUD4 outside the DUB domain can switch Ub chain cleavage specificity from K48 to K63 linkages^32^. When the cleavage reaction was allowed to proceed longer, OTUD4 was able to cleave other Ub chain linkages (K6-, K11-, K27-, K29-, and K33-). Comparison of WT OTUD4 with the G398V mutant showed that the cleavage of all Ub chain types were reduced when the G398V mutant was used, except for K63-linked Ub chains that showed compatible activities with WT OTUD4 (Figure 4D). These findings imply that the OTUD4 GHS mutant has tempered activity toward various Ub linkage types, except for K63-linkages.

We next determined the activity of OTUD4 in cleaving various Ub chain types assembled by RNF216. Results using TUBE assays indicated that expression of WT OTUD4 in HEK293 cells reduced K63-linked Ub chains on RNF216. Based on these results, single Lys mutants of Ub were co-expressed with RNF216 (Figure 4E and Figure S5D). As expected, RNF216 had a strong bias to assemble K63-Ub chains. However, we also found that RNF216 could assemble other chains with the following hierarchy: K63 > K48 > K6 > K11 > K27. Notably, the DUB domain of OTUD4 could de-ubiquitinate all chain types on RNF216 with modest effects on K27-linked Ub chains (Figure 4E). These findings indicate that in addition to K63, OTUD4 has the capacity to cleave multiple Ub chain types on RNF216.

To further glean insight into functions of OTUD4 and its role in GHS, we characterized the *Otud4* KO mouse (Figure S6A)^32^. *Otud4* KO male mice had reductions in testicular size indicating reduced breeding viability (Figure S6B), which phenocopied our findings in *Rnf216* KO mice^24,25^. Since OTUD4 is a DUB for RNF216, we next evaluated its biological significance. To do this, we mirrored our ubiquitination and steady-state assays for OTUD4 in the *Rnf216* KO background but instead monitored RNF216 in an *Otud4* KO background. First, we generated *Otud4* GT1-7 KO cells using the Crispr-Cas9 method (Figure S6C). RNF216 levels were not significantly altered in *Otud4* KO cells as shown by the anisomycin pulse-chase assay (Figure 5A). Surprisingly, removal of *Otud4* decreased RNF216 ubiquitination in GT1-7 cells and *Otud4* KO mouse brain (Figure 5B and 5C). Steady-state levels of RNF216 were also not affected in *Otud4* KO GT1-7 cells (Figure 5D) and *Otud4* KO mouse hypothalamus (Figure 5E). Taken together, while these findings show a connection of OTUD4 as a DUB for RNF216, OTUD4 does not directly alter RNF216 stability in the cell.

**Figure 5:**
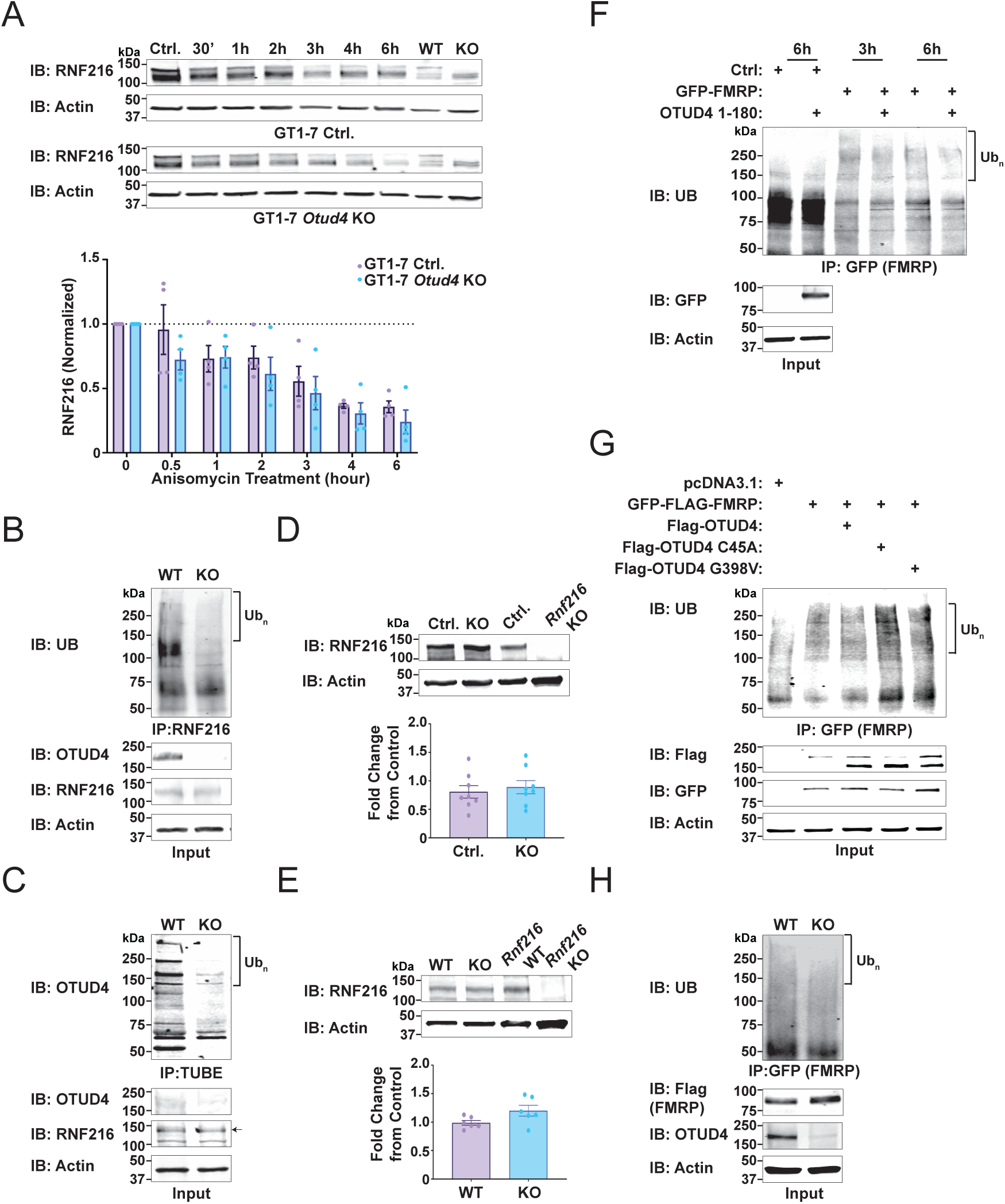
OTUD4 deubiquitination of RNF216 is not destabilizing and the translational repressor FMRP is an OTUD4/RNF216 shared substrate. (A) *Top*, RNF216 half-life is unchanged in *Otud4* KO GT1-7 cells. Crispr control (WT) and *Otud4* KO (KO) GT1-7 cells were treated with vehicle (DMSO) or anisomycin (20 µM) for 30’, 1h, 2h, 3h, 4h, or 6h. Processed samples were subjected to immunoblotting with anti-RNF216 and -Actin antibodies. *Bottom*, Quantification of RNF216 in GT1-7 Control and GT1-7 *Otud4* KO cells. One-way ANOVA with Dunnett’s multiple comparisons test. N = 4 independent biological replicates. (B) RNF216 ubiquitination is reduced in *Otud4* KO GT1-7 cells. *Otud4* GT1-7 WT and KO cell lysates were immunopurified under denaturing conditions with an anti-RNF216 antibody. Processed samples were immunoblotted with an anti-UB antibody. Inputs were immunoblotted with anti-RNF216, -OTUD4, and -Actin antibodies. N = 3 independent biological replicates. (C) RNF216 ubiquitination is also reduced in *Otud4* KO whole brain. *Otud4* WT and KO whole brain tissue lysates were evaluated for changes in RNF216 ubiquitination using TUBEs and processed samples were immunoblotted with an anti-RNF216 antibody. Inputs were immunoblotted as in B. (D) *Top*, No change in steady state RNF216 in *Otud4* KO GT1-7 cells. *Bottom*, Quantification of RNF216 levels normalized to Actin. N = 8 independent biological replicates. (E) *Top*, No significant change in RNF216 in P40 *Otud4* KO mouse brain. *Bottom*, Quantification of RNF216 levels normalized to Actin. Welch’s t test, P = 0.056. N = 6 independent biological replicates. (F) OTUD4 deubiquitinates FMRP in vitro. HEK 293 cells were transfected with GFP-FMRP. Cell lysates were immune purified under native conditions using an anti-GFP antibody as stated in the methods and then incubated with recombinant OTUD4 (1-180). Terminated reactions were immunoblotted with an anti-UB antibody. (G) OTUD4 deubiquitinates FMRP in cells, which is disrupted upon mutation of its catalytic domain and in the GHS mutant. HEK 293 cells were transfected with Flag-OTUD4, Flag-OTUD4 C45A, and FLAG-OTUD4 G398V with and without GFP-FMRP. Cell lysates were immunopurified under denaturing conditions using an anti-GFP antibody and processed samples were immunoblotted with an anti-UB antibody. Inputs were immunoblotted with anti-Flag, -GFP, or -Actin antibodies. (H) FMRP ubiquitination is reduced in *Otud4* knockout cells. GFP-FMRP was transfected in Ctrl. or *Otud4* KO GT1-7 cells. Cell lysates were immunopurified under denaturing conditions using an anti-GFP antibody and processed samples were immunoblotted with an anti-UB antibody. Inputs were immunoblotted with anti-Flag, -OTUD4, or -Actin antibodies.

### FMRP is a shared substrate of RNF216 and OTUD4

To understand RNF216 relationships with OTUD4, we surveyed the OTUD4 interaction network and found common proteins that were classified as RNF216 substrates, with many of those targets known to be mutated in neurological disease (Figure S6D, red). When evaluating their biological significance, shared targets were also involved in mRNA processing, DNA damage, and translation regulation (Figure S6E), implying a shared functional nexus between these two enzymes. One shared target was FMRP, an established neuronal translational repressor. In vitro, addition of recombinant OTUD4 was able to decrease FMRP ubiquitination (Figure 5F). When expressed in cells, OTUD4 modestly reduced the ubiquitination of FMRP, which required the catalytic domain of OTUD4 (C45A). Moreover, the OTUD4 GHS mutant (G398V) was less effective than WT OTUD4 in removing Ub chains conjugated to FMRP (Figure 5G). Like *Rnf216* KO reductions on OTUD4 ubiquitination (Figures 2A and 2B), a loss of OTUD4 paradoxically led to a decrease in ubiquitinated FMRP in GT1-7 cells (Figure 5H). Taken together, our results show that FMRP is an RNF216 and OTUD4 substrate, but it seems that removal of either of these enzymes produces synonymous effects on the ubiquitination of their common substrates in cells. Given this finding, we propose that RNF216 and OTUD4 form a catalytic cycle to tune protein ubiquitination of shared substrates.

### RNF216 and OTUD4 regulate protein synthesis in cells

Given the finding of FMRP as a substrate for RNF216, shared interaction partner for OTUD4, and common functions of RNF216 and OTUD4 regulating protein synthesis and DNA repair, we further probed the biological significance of RNF216-targeted substrates. To do this, we performed the RNF216 OUT assay in HEK293 cells (Figure S7C) and compared the substrate profile to substrates identified in GT1-7 cells (Figure 6A-C). Out of 174 GT1-7 and 62 HEK293 substrates, there were only 3 that were overlapping (MYH9, PARP1, and RPS13) (Figures 6A, 6C). When evaluating the separate pool of cell substrates based on their biological significance, the most significant functions were related to protein translation and DNA repair (Figure 6D, Figure S8).

**Figure 6:**
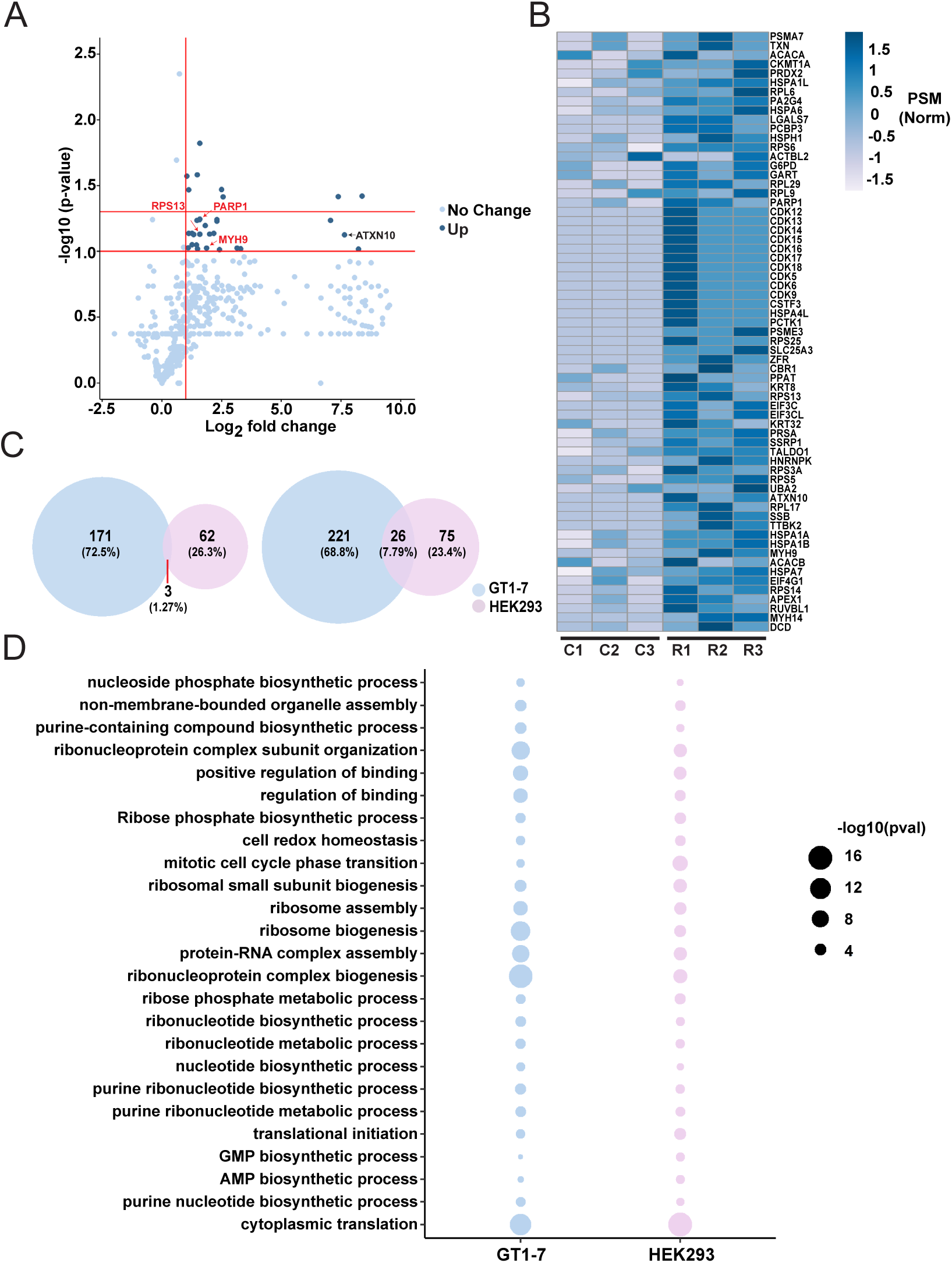
RNF216 substrate pools differ between cell types but have shared biological functions related to protein synthesis. (A) Volcano plot of RNF216 substrates identified from OUT in HEK293 cells. The dark blue dots show the differential expression of targets in xRNF216 compared to xRNF216 CA, Log 2 [PSM ratio xRNF216/ xRNF216 with CA mutation] > 1 and -Log 10 p > 1. N = 3 independent biological replicates. (B) Heatmap for significant targets as calculated using normalized PSM values from OUT. C1-C3: 3 replicates from xRNF216 CA control cells. R1-R3: 3 replicates from xRNF216 cells. (C) *Left*, Comparison of substrates of RNF216 from OUT in GT1-7 and HEK cells. Substrate overlap of RNF216 substrates between HEK 293 and GT1-7 cells reveals only 3 that are common. *Right*, Functional annotation of shared biological functions between HEK 293 and GT1-7 cells. Functional annotation was performed using Clusterprofiler4.0. (D) The most significant shared biological function of RNF216 substrates between HEK 293 and GT1-7 cells is cytoplasmic translation.

Given our bioinformatics findings and validation of the translational repressor FMRP as a target, we next sought to evaluate if RNF216 could regulate protein synthesis. Protein synthesis was measured across nonneuronal and neuronal systems upon manipulating RNF216 using puromycylation assays^38,39^. Transfection of RNF216 in HEK293 cells led to an increase in protein synthesis (Figure 7A). Given the strong link between dendritic protein synthesis and dendrite development, we next looked at the effect of RNF216 manipulations on dendritic morphology in developing hippocampal neurons. Deletion of *Rnf216* reduced dendritic complexity in developing primary hippocampal neurons (Figure 7B). Taken together, our findings show that while RNF216 promotes the ubiquitination and turnover of proteins, it also regulates substrates that modulate protein synthesis. Moreover, our findings imply that alterations in RNF216 in GHS not only decrease protein degradation but also reduce protein synthesis.

**Figure 7:**
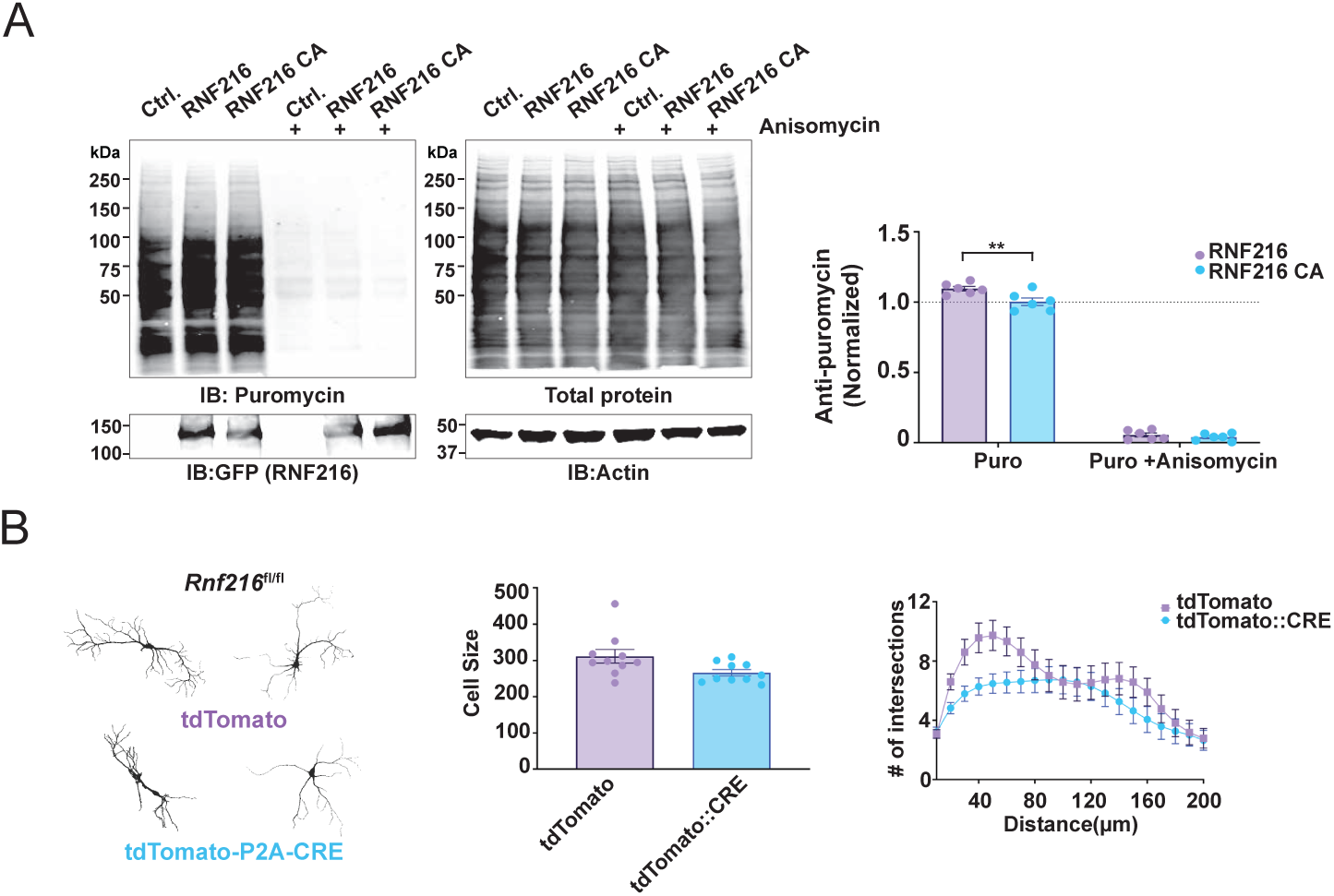
RNF216 is involved in the regulation of protein synthesis in cells. (A) *Left*, RNF216 overexpression increases protein synthesis in HEK 293 cells. HEK 293 cells were transfected with GFP-RNF216 WT or control. Cells were then treated with 10 µM puromycin or 10 µM PMY and 20 µM Anisomycin for a duration of 30 minutes. Processed cell lysates were then immunoblotted with an anti-puromycin and total protein stain followed by-GFP, or -Actin antibodies. *Right*, Quantification of puromycin labeling normalized to total protein stain. N = 6 independent biological replicates. Two way ANOVA, Sidak’s multiple comparison P = 0.0063. (B) Conditional deletion of *Rnf216* reduces cell size and dendritic complexity in developing primary hippocampal neurons. *Rnf216^fl/fl^* primary hippocampal neurons were transfected at DIV3 with plasmids expressing CamKII-tdTomato control or CamKII-tdTomato-P2A-CRE. Neurons were fixed at DIV15, imaged and then quantified using Sholl analysis as described in the methods. *Left*, Representative images of traced neurons from each experimental group. *Middle*, Quantification of cell size. Unpaired t-test p = 0.0405. *Right*, Sholl analysis demonstrating a decrease in the number of dendritic branch crossings in CRE expressing *Rnf216^fl/fl^* neurons. Two way ANOVA P < 0.0001

Given this unexpected finding with RNF216, we were compelled to consider that other Ub E3 ligase enzymes could serve a similar role. To support this idea, we analyzed shared biological functions of substrates for 5 different E3 Ub ligases identified by OUT in HEK293 cells (UBE3A^40^, RNF38^41^, PARKIN^29^, RNF216, and CHIP^42^). Strikingly, 4 out of 5 of these E3 Ub ligases shared a common biological function of translation initiation (Figure S9A), representing HECT, RBR, and U-Box classes of E3s. This also included CHIP, another E3 ligase mutated in GHS (Figures S9B and S9C)^9^. Taken together, our findings indicate that in addition to RNF216, specific classes of E3s may also regulate protein synthesis.

## Discussion

In this study, we generated an OUT system for RNF216, a RBR E3 Ub ligase that is mutated in GHS. Using OUT, we identified 174 substrates in GT1-7 hypothalamic cells and 65 in HEK293 cells. From the hypothalamic substrate list, the DUB OTUD4 stood out, as compound heterozygous mutations in *OTUD4* have also been reported in GHS^8^. Since RNF216, as an E3, could promote OTUD4 ubiquitination leading to its instability, we proposed OTUD4 and RNF216 would antagonize each other with RNF216 serving a regulatory role, tempering OTUD4 DUB activity to bias ubiquitination of RNF216 substrates. We also proposed that OTUD4 was a DUB for RNF216 substrates. To support this, we overlaid the RNF216 substrate network identified by OUT with the OTUD4 interaction network and found 32 common targets. Many of these targets, which include the translational repressor FMRP and another translational regulator ATXN2 are known to be mutated in other neurological disorders. ATXN2 is highly significant given mutations result in Spinocerebellar Ataxia Type 2 (SCA2), a cerebellar ataxic neurodegenerative disorder that has similar phenotypes to GHS^43^.

Using in vitro and cell-based assays, we found that OTUD4 could deubiquitinate the RNF216 substrate FMRP. FMRP is known to exist in complex with OTUD4 at RNA granules in neuronal dendrites^44^, which are formed upon environmental stress and are hotspots for regulating protein translation^45,46^. Upon evaluation of RNF216 ubiquitination in a spatial context, the addition of WT but not catalytic inactive RNF216 increased the formation of ubiquitinated OTUD4 (Ub-OTUD4) complexes in the soma and dendrites of primary hippocampal neurons whereas deletion of RNF216 reduced the formation of these complexes. Furthermore, RNF216 facilitated the formation of Ub-OTUD4 donut-like structures in the soma and dendrites of neurons. While out of scope for this study, the significance and identity of these structures will be an active area of future research.

Using in vitro and cellular assays, RNF216 promoted the ubiquitination of OTUD4, which was enhanced when the catalytic activity of OTUD4 was blocked by an active site mutation (C45A). RNF216-mediated ubiquitination of the GHS-associated mutation (G398V) was also reduced in cells. On the other hand, OTUD4 removed Ub chains conjugated to RNF216 in vitro and in cells, and the G398V mutant of OTUD4 had tempered activity toward cleaving Ub chains and exhibited modest reductions in its ability to deubiquitinate FMRP when overexpressed. While the OTUD4 GHS mutant had reduced DUB activity, we did not see dramatic effects on its ability to remove RNF216 assembled Ub chains. One explanation for this perplexing finding is that the OTUD4 GHS mutant may have selective deficits in cleaving Ub chain types, which is supported by our in vitro assays demonstrating that this mutant has reduced activity in cleaving a variety of di-Ub linkages with no effects on K63-. Given our findings that RNF216-dependent Ub assembly on OTUD4 is K6-biased, the reduction could be related to a feedback loop or an inability to assemble these chain types. In contrast, our findings show that OTUD4 dominantly disassembles K63-linked Ub chains on RNF216. This finding in combination with our in-vitro results on a lack of effect on K63-Ub cleavage favors the model where the OTUD4 GHS mutation may have a selective deficit in the cleavage of Ub chain types.

The potential regulatory feedback loop between OTUD4 and RNF216 using in vitro and cell-based assays shows that OTUD4 could deubiquitinate RNF216. RNF216 and OTUD4 are also colocalized in neurons. These findings and descriptions above support a functional connection for these enzymes to work as a pair. Evidence for DUB and E3 ligase pairing has been supported by findings that DUBs have E3 binding partners and with examples of DUBs that alter E3 Ub ligase activity^47,48^. While most studies show that DUBs dominantly exert actions to facilitate E3 activity by regulating their stability, we found that RNF216 stability was unaffected in *Otud4* KO GT1-7 cells. In our study, OTUD4 removes RNF216 assembled Ub chains of various linkage types, with a high reactivity towards K63-linkages. Thus, we posit that OTUD4 could operate to remove inhibitory RNF216 chains to facilitate RNF216 activation. Alternatively, OTUD4 Ub chain editing may be an essential step for the full-scale assembly of ubiquitinated RNF216 substrates, especially given known roles of RNF216 in heterotypic Ub chain assembly^17,19^. This could explain why depleting OTUD4 decreases ubiquitination of the RNF216 substrate FMRP. Importantly, mutations in RNF216 or CHIP have been reported in GHS^8,9^. The operational rules of E3s and their role in GHS may be a broader theme as another study found the DUB USP7 was a binding partner for the E3 Ub ligase CHIP^34^. Here, Tau was identified as a CHIP and USP7 substrate, but this pairing operated in an antagonistic manner as they competed for a similar Tau interaction site. While USP7 could deubiquitinate CHIP, CHIP did not alter USP7 ubiquitination. While this study describes a unidirectional regulation of E3-DUB pairing, it highlights the importance in their capacity to engage with E3 substrates, which has strong implications for GHS.

Given our results supporting RNF216-OTUD4 interactions, we sought to provide a model for how they could work with or against one another. While we initially thought that OTUD4 could counteract RNF216 assembly to compete and remove Ub chains, this did not appear to be the case, as deletion of *Otud4* in cells also reduced the ubiquitination of RNF216 substrates such as FMRP as well as RNF216. Therefore, we propose a model where an RNF216-OTUD4 pair functions to add, trim, and edit RNF216 assembled chains on its substrates. While preliminary, this model is partially supported by the finding that deletion of *Otud4* in mice results in a similar phenotype as *Rnf216* removal that includes reductions in male testicular size, a prominent feature of GHS^8,24,49^ and the ability of RNF216 to assemble a variety of Ub chain linkages that also include formation of heterotypic chains^17,19,33,50^. There must be factors that can facilitate the formation of these atypical linkages. The introduction of a DUB to support chain editing and trimming of RNF216 substrates would produce the variety of RNF216 Ub configurations that can assemble on to its substrates. OTUD4 is a highly disordered protein but does have a characterized Ub binding domain lying adjacent to its catalytic OTU region that could serve as a transient interaction site to sculpt heterotypic chain assembly^32^. Importantly, the diversity of RNF216 chains could complement engagements with different DUBs. OTUB1 was another DUB identified in the RNF216 OUT screen so it is likely that this DUB-E3 pairing could manage the ubiquitination of other RNF216 substrates that were not identified in the OTUD4 network (e.g. Tau^19^).

In the developing brain, rates of protein synthesis slightly outweigh rates of protein degradation^51–53^. However, post development, the balance of protein synthesis and degradation are necessary to maintain protein homeostasis^51^, which is governed by numerous adapters and large molecular machines (e.g. ribosomes and the proteasomes)^54^. Disruption of protein synthesis and degradation occurs in pathological conditions that span the spectrum of cancer, development, mental health, and neurodegeneration^54,55^. While factors within the proteostasis network have been characterized^56^, the molecular determinants that maintain proteostasis are not well established. In this study, we unexpectedly found that changes in RNF216 expression not only regulate ubiquitination and stability of its substrates but also had effects on protein synthesis. The translational repressor FMRP as an RNF216 substrate provided the first line of evidence that its function could be to positively regulate protein synthesis rates. Disruption in the balance of protein synthesis and degradation are thought to drive neurological disease^54,55^, and recent studies have shown that use of inhibitors to block proteasome-dependent degradation also reduces protein synthesis in neurons^57^. From a general standpoint, the loss of proteins via their degradation must have a compensatory feedback loop to make new proteins to replace those that are lost, similar to how neurons are under homeostatic control to maintain set firing rates which is necessary for stabilizing neural circuits^58^. Given this unexpected role for RNF216, we propose that it likely serves as a molecular “proteostat”. The definition of a molecular proteostat must fulfill a series of criteria that encompass classifications of proteins that make up the proteostatic network^56,59,60^ that include 1) An ability to monitor translation rates; 2) An ability to regulate chaperone substrates; 3) The ability to target proteins to degradation systems and ; 4) The ability to regulate stress response proteins. We believe that RNF216 fulfills these four criteria given its known existing functions and identification of new substrates found here.

While RNF216 fulfills the criteria of a “proteostat”, additional E3 Ub ligases may subserve a similar role. To support this notion, we compared all substrate profiles identified using the OUT platform in HEK293 cells (UBE3A^40^, RNF38^41^, PARKIN^29^, RNF216, and CHIP^42^). E3 ligase combinations had some overlapping substrates as well as shared biological functions. However, when searching for a common biological function across all E3 enzymes, 4 out of 5 had a shared function of “translation initiation”. This included classes of E3s falling into RBR, HECT, and U-Box-containing E3s. The notable exception was RNF38, which is a single monomeric nuclear enriched RING E3^61^. While RNF38 is a poorly characterized substrate, we recently found that this E3 regulates nuclear import-export movement of proteins so its function may be nondegradative^41^ as opposed to the other E3s, which have well established degradative functions in cellular systems. Future directions should target generating additional OUT platforms for other E3s for direct proteomic comparison and should formally test the role of other E3 ligases in regulating protein synthesis. Our findings also lend a cautionary tale of traditional E3 ligase capture approaches identifying differentially expressed ubiquitinated proteins in proteomic platforms without controlling for gross changes in protein synthesis. Use of traditional methods^62^ would require a multi-omics approach, which disentangles changes in protein synthesis rates to alterations in posttranslational Ub modification status.

Taken together, we propose that RNF216 is a molecular proteostat, whose function is to regulate a tailored suite of substrates that maintain the proteostatic network. It is likely that many E3 ligases share this feature, having specialized protein ensembles that can maintain the proteostatic landscape. These findings could explain why mutations in E3 Ub ligases such as RNF216 are pervasively found to be mutated and disrupted in numerous neurological conditions^6,63^.

## Limitation of the study

This study takes an overexpression orthogonal approach to generate point mutations in protein ubiquitination enzymes that may change the Ub conjugation topology on endogenous substrates. While OUT has the advantage of direct transfer to endogenous protein targets, secondary validation of substrates and mapping of Ub chain topology must be performed using WT enzymes. Furthermore, the introduction of orthogonal mutations may alter E3-substrate interfaces, making it difficult to capture the entire repertoire of E3 ligase substrates. Regardless of these caveats, the example provided here demonstrates that the RNF216 OUT platform can identify important substrates that can uncover new biological mechanisms and provide new insights into the pathology of GHS.

## Resource availability

### Lead contact

Further information and requests should be directed and will be fulfilled by the lead contact, Angela Mabb

### Materials availability

All reagents generated in this study are available from the lead contact.

### Data and code availability

## Acknowledgements

This work was funded by NIH/NINDS R01NS136464, NIH/NINDS R21NS116760, and a Brains & Behavior Seed Grant to A.M.M. and J.Y.; National Science Foundation CAREER Award (2047700) to A.M.M.; R01 GM160854, R01 CA282733, and P01 CA092584 to N.M. Imaging research reported in this publication was supported by the Office of The Director, National Institutes Of Health and National Institute Of General Medical Sciences under Award Number S10OD032336-01. The content is solely the responsibility of the authors and does not necessarily represent the official views of the National Institutes of Health.

## Author contributions

Conceptualization, A.M.M. and J.Y.; methodology, W.W., R.L., J.Z., S.L., A.J.C., D.A., Z.D.A., D.W., E.K.; investigation, all authors; writing, W.W., J.Z., S.L., D.A., Z.D.A., N.M., J.Y., A.M.M.; funding acquisition, A.M.M. and J.Y.; resources, A.M.M., J.Y., and N.M.; supervision, A.M.M. and J.Y.

## Declaration of interests

The authors declare no competing interests.

## RESOURCE AVAILABLITY

### Lead contact

Further information and requests for resources and reagents should be directed to and will be fulfilled by the lead contact, Dr. Angela Mabb.

### Materials availability

The availability of GT1-7 CRISPR lines requires initiation of a material transfer agreement (MTA) with the Salk Institute for Biological Sciences.

Plasmids generated in this study will be shared by the lead contact upon request.

All mouse lines generated in this study are commercially available. *Otud4* knockout mouse requests can be directed to the lead contact or Dr. Nima Mosammaparast.

### Data and code availability

All data reported in this paper will be shared by the lead contact upon request. Any additional information required to reanalyze data reported in this paper is available from the lead contact upon request.

## EXPERIMENTAL MODEL AND SUBJECT DETAILS

### Animals

All animals were kept in standard housing with littermates, provided with food and water ad libitum and maintained on a 12:12 (light-dark) cycle and carried out in accordance with the National Institutes of Health Guidelines for the Use of Animals using approved protocols by the Georgia State University Institutional Animal Care and Use Committee. Animals used in primary hippocampal cultures are balanced for sex and littermates from heterozygous pairings. *Otud4* knockout mice were generously provided by Dr. Nima Mosammaparast^32^.

### Cell cultures and transfections

HEK293 cells were generously provided by Dr. Jun Yin (Georgia State University). Cells were maintained in Dulbecco’s Modified Eagle Medium (DMEM; Corning # 10013CV) containing with 10% FBS and 1% penicillin-streptomycin (ThermoFisher). 2x10^6^ cells were transferred to a 10cm dish one day prior to transfection with Lipofectamine 2000 (ThermoFisher) according to the manufacturer recommendations. Cells were harvested after incubation at 37°C for 24 hours. To control for transfection efficiency, the plasmid pcDNA3.1 was added to maintain the identical DNA concentrations with other conditions that had multiple DNAs that were transfected.

GT1-7 cells were generously provided by Dr. Pamela Mellon^64^. Cells were maintained in DMEM with 10% FBS and 1% Pen Strep at 37°C with 5%CO2. For proteasome activity experiments, cells were seeded at a density of 2.5 x 10^5^ cells/well in a 6-well dish. Then cells were treated with 10µM MG-132 (sigma 474790) for 4 hours before harvesting. For OUT system expression, 2x10^6^ cells were transferred to 10cm dishes one day prior to transfection with Lipofectamine 3000 (ThermoFisher) according to the manufacturer recommendations. Cells were harvested after incubation at 37°C for 24 hours.

### Primary Hippocampal Neuronal Cultures

Primary hippocampal neurons of mixed sex were isolated from P0-1 mice as previously described^65^ and cultured on poly-d-lysine-coated coverslips (0.1 mg/ml) in 24-well plates at a density of 75,000 cells/well or in 6-well plates at a density of 100,000 cells/well. Cultures were maintained in neuronal feeding media: Neurobasal media (Gibco) containing 1% GlutaMAX (Gibco), 2% B-27 (Gibco), 4.8 μg/mL 5-Fluoro-2’-deoxyuridine (Sigma), and 0.2 μg/mL Gentamicin (Sigma). Cultures were transfected with Lipofectamine 2000 on DIV 10 as described in ^50^ with equal amount of cDNA transfected into all conditions. GFP or Td-tomato were used as a cell fill to identify neuron morphology as described in^66,67^.

### Plasmid Construction

Flag-OTUD4, Flag-OTUD4C45A were gifts from Nima Mosammaparast^32^. Flag-OTUD4-G398V mutant was generated by PCR-based mutagenesis using Flag-OTUD4 as a template. pCAGGS-xFlag-UBA1-P2A-xMyc-RNF216 WT and pAAV-xV5-UBCH7-P2A-HBT-xUB were generated using a two-step cloning process. First, the xFlag-UBA1-P2A-xMyc-RNF216 and xV5-UBCH7-P2A-HBT-xUB inserts were synthesized with embedded restriction sites integrated into the pcDNA3.1(+) vector backbone (Genscript). pcDNA3.1(+) xFlag-UBA1-P2A-xMyc-RNF216 was digested with KpnI and NheI restriction enzymes and subcloned into the pCAGGS-plasmid backbone (Addgene 87221). pcDNA3.1(+) xV5-UBCH7-P2A-HBT-xUB was digested with BamHI and EcoRI restriction enzymes and subcloned into the pAAV-CAG- (Addgene 59462) plasmid backbone. pCAGGS-xFlag-UBA1-xMyc-RNF216 CA was generated by PCR-based mutagenesis using pCAGGS-xFlag-UBA1-P2A-xMyc-RNF216 as a template.

### Generation of control and OTUD4 Crispr clones and validation of Crispr sequences

Two sgRNA targets (*Otud4-1* and *Otud4-2*) in the pLentiCRISPRv2 backbone along with custom sequencing primers targeting mouse *Otud4* were obtained from Genscript. GT1-7 cells were maintained as previously described^24^. Cells were seeded at a density of 2.5x10^5^ cells/well in a 6-well dish and transfected with CRISPR-Cas9 plasmids using the standard Lipofectamine 3000 (Thermo Fisher) protocol. Briefly, 2.5 μg of plasmid DNA was mixed with Lipofectamine 3000 reagents and incubated for 4 h before replacing with fresh prewarmed media. After 72 h, 2.0 μg/mL of puromycin was added to the media for selection. After integration was established, single cell clones were selected and expanded into colonies under puromycin selection. Loss of OTUD4 was validated through immunoblotting. CRISPR clones that showed a loss of OTUD4 were used for validation and downstream experiments. Out of the 2 targets, Otud4-2 was the only target that generated OTUD4 deficient clones.

OTUD4 deficient GT1-7 clones were plated at a density of 1.0 x10^6^ in a 10cm dish. Genomic DNA was extracted using the DNeasy Blood & Tissue Kit (QIAGEN) according to the manufacturer’s protocol. The concentration and purity of purified DNA was assessed using a NanoDrop ND-2000/2000c (ThermoFisher). DNA was only used if it had an A260/A280 ratio between 1.8–2.0. Purified PCR products were submitted to GENEWIZ (South Plainfield, NJ) for sequencing.

### Purification of recombinant OTUD4 and RNF216

Recombinant pET plasmids for the expression of RNF216 and OTUD4 1-180 were transformed into BL21 cells and cultured in 2×YT broth with antibiotics under 37°C until the OD values of the cultures were in the range of 0.6–0.8. IPTG was added to the cell culture to reach the final concentration of 1 mM to induce protein expression, and the cell culture was incubated overnight under 16°C with agitation at 220 round per minute (rpm) before the cells were harvested by centrifugation (5,500 rpm, 4 °C, 20 min). 50 μM ZnCl2 was additionally added to enhance the expression of RNF216. Cells were resuspended in 20 mL of lysis buffer (8 mM NaH2PO4, 300 mM NaCl, 10 mM imidazole, pH 8.0) with the addition of 1mM PMSF, and the suspension was incubated on ice for 30 min. The cells were lysed by sonication on ice and the cell lysates were centrifuged (10,000 rpm, 4°C, 60 min). After centrifugation, the supernatant of the lysate was collected to bind with Ni-NTA beads. The protein was purified by a gravity flow column with washes of 30 mL lysis buffer (50 mM Tris, 300 mM NaCl, 5 mM imidazole, pH 8.0) once and 30 mL wash buffer (50 mM Tris, 300 mM NaCl, 20 mM imidazole, pH 8.0) twice followed by elution with 5 mL of elution buffer (50 mM Tris, 300 mM NaCl, 250 mM imidazole, pH 8.0). The eluted protein solution was further dialyzed against a dialysis buffer (50 mM Tris, 50 mM NaCl, 1 mM DTT, pH 8.0) and concentrated. The final protein solution was aliquoted and stored at –80℃.

### Western Blotting

HEK293 or GT1-7 cells were harvested and then cell pellets were lysed on ice in radioimmunoprecipitation assay (RIPA) buffer (150 mM NaCl, 50 mM Tris-HCl, 1% v/v Nonidet P-40, 0.5% sodium deoxycholate, 0.1% SDS) with 1 mM DTT (Millipore) and protease inhibitors (0.1 mM PMSF (Calbiochem), 1 μM leupeptin (Millipore), 0.15 μM aprotinin (Millipore). Lysates were centrifuged at 13,000 rpm for 25 min at 4°C to precipitate insoluble extracts. Protein concentrations were measured using the Pierce 660 assay (ThermoFisher).

Cell lysates were heated for 5-7min at 45°C prior to load them on a 4–15% gradient with 4X Protein sample loading buffer (LI-COR) containing 0.001% β-mercaptoethanol. Proteins were then transferred to nitrocellulose membrane at 4°C. Membranes were blocked overnight at 4° in Intercept tris-buffered saline (TBS) blocking buffer (LI-COR) then probed in primary antibodies in 1:1 blocking buffer to 1% Tween-20 (Acros) in TBS (TBST) with 0.02% NaN_3_ overnight at 4°C. Membranes were washed 3 times with de-ionized water (DI water) for 5 min, and incubated with secondary antibodies in 1:1 blocking buffer to TBST 0.1% SDS (Bio-Rad) were added to the membranes for 1 hr at room temperature, then washed 2 times with TBST and 1 time with DI water for 5 min.

Western blot membranes were scanned using the LI-COR Odyssey CLx scanner (LI-COR Biosciences). Images were analyzed using ImageJ software (NIH) with the Gel Analysis tool.

### Ubiquitin Assays

HEK293 or GT1-7 cells were transfected with DNA plasmids 24 h and harvested and lysed in RIPA buffer 24 hrs later. Collected supernatants from samples were diluted to with RIPA buffer, which contained protease and phosphatase inhibitors as above. Expressed protein was incubated with primary antibody for 1h at 4°C then an equal volume of pre-equilibrated Protein A/G PLUS-Agarose Beads were added to each sample and left to tumble overnight at 4°C. The supernatant was discarded and beads were washed with RIPA buffer three times and heated in 4X Protein sample loading buffer and separated by SDS-PAGE.

Tandem Ubiquitin Binding Entities (TUBEs) were used according to the manufacturer’s protocol. Cells or brain tissue were lysed in buffer containing protease and phosphatase inhibitors as required by the protocol. Lysates were centrifuged at 13,000 rpm for 25mins at 4°C. After measuring the protein concentration, 1mg of protein was taken for each sample and brought up to a total of 1ml in lysis buffer. Agarose beads were pre-equilibrated in buffer with inhibitors. Protein samples were tumbled with agarose at 4°C overnight. Following the same washing and protein separation step, membrane was incubated in primary Ab.

Reconstituted ubiquitination reactions were set up in 50 μL of reaction buffer (50 mM Tris, 5 mM MgCl2, 5 mM ATP, and 1 mM DTT, pH 7.5). WT full-length human RNF216 protein (3 μM) was incubated with 1 μM WT Uba1, 5μM WT UbcH7, 20 μM WT HA-UB at 37 °C for 4 hours. Parallel control reactions were also set up with each of the enzyme components excluded from the reaction mixture. The reactions were quenched by 3U apyrase at room temperature for 10 minutes. Then the reaction mixtures were incubated with purified recombinant OTUD4 (1-180) for 6 hours. The reaction was quenched by boiling the samples in the SDS-PAGE loading dye with DTT for 5 min and analyzed by SDS-PAGE and western blot probed with the RNF216-specific antibodies.

#### Tetra ub assay

0.4 ug Tetra-ub was treated with 10uM recombinant OTUD4 (1-180) in PBS buffer. The reaction was quenched by boiling the samples in the SDS-PAGE loading dye with DTT for 5 min and analyzed by SDS-PAGE and western blot probed with the Ub antibodies.

### Immunocytochemistry

48 h after transfection, neurons were fixed for 20 min in cold PBS containing 4% paraformaldehyde/4% sucrose solution at 4°C. After washing with cold PBS 3 times, neurons were permeabilized with 0.2 % Saponin for 15 min then blocked in 10 % normal goat serum (NGS) in PBS for 1 hr at 37°C. Neurons were then incubated overnight at 4°C in primary antibody in 3% NGS. On the 2^nd^ day, neurons were washed 3 times in 3% (vol/vol) NGS in PBS and then incubated in secondary antibody at 1:1,000 in 3% NGS for 1 h in the dark at room temperature. Coverslips were washed with phosphate buffer saline (PBS) then mounted onto slides with Fluorogel (Electron Microscopy Sciences).

### Puromycylation

48 h after transfection, 100 μM puromycin or 100 μM puromycin and 200 μM emetine^39^ incubate at 37°C for 5min, then neurons were placed on the ice and washed by cold PBS with 0.0003% digitonin. After washing, neurons were fixed for 20 min in cold PBS containing 4% paraformaldehyde and 4% sucrose solution at 4°C. After washing with cold PBS 3 times, neurons were permeabilized with 0.2 % Saponin for 15 min then blocked in 10 % NGS in PBS for 1 hr at 37°C. Neurons were then incubated overnight at 4°C in primary antibody in 3% NGS overnight. On the 2^nd^ day, neurons were washed 3 times in 3% (vol/vol) NGS in PBS and then incubated in secondary antibody at 1:1,000 in 3% GHS for 1 h in the dark at room temperature. Coverslips were washed with phosphate buffer saline (PBS) then mounted onto slides with Fluorogel (Electron Microscopy Sciences).

For HEK293 and GT 1-7 cell lines, cells were treated with 10μM puromycin or 10μM puromycin with 20μM Anisomycin separately for 30 mins then harvested and the cell pellet was lysed on ice in RIPA buffer. Samples were heated in 4X Protein sample loading buffer and resolved by SDS-PAGE

### Proximity ligation assay (PLA)

48 h after transfection, cells were fixed for 20 min in cold PBS containing 4% paraformaldehyde/4% sucrose solution at 4°C. After washing with cold PBS 3 times, neurons were permeabilized with 0.2 % Saponin for 15 min then blocked in 10 % NGS in PBS for 1 hr at 37°C. Neurons were then incubated overnight at 4°C in primary antibody in 3% NGS. The primary Ab was washed off using the washing buffer from Millipore Sigma (Duo 92008), incubated with PLA probe at a 1:5 dilution for 1 hr at 37 °C. The ligase with ligation buffer was then added at a 1:40 dilution and incubated for 30mis at 37°C. After washing, the signal was amplified by adding 1:80 Polymerase for 100 mins at 37°C. The amplification solution was removed by washing the slide using buffer B and coverslips were mounted on microscope slides.

### Image Acquisition and Analysis

For primary hippocampal neurons, images were acquired using a LSM 700 (Zeiss) or LSM 980 with Airyscan confocal microscope under 40x objective. Only cells that had a distinctive pyramidal morphology were imaged. Images were acquired with identical acquisition parameters (gain, contrast, pinhole) for each marker per imaging session and treatment conditions were interleaved during each imaging session. Images were analyzed using ImageJ software (NIH). For characterization of Ub-OTUD4 “donut” structures, the region of interest was selected based on the neuron morphology outlined by the GFP fluorescent signal in neuronal soma and dendrites. Based on the PLA fluorescence pattern, cells were manually subdivided into three categories: donut (ring with a black dot in the center), spherical, and intermediate-shaped (in between donut and spherical shapes). The average puncta area in the soma and dendritic regions were measured in each neuron and the experimenter was blinded to treatment conditions before imaging and then unblinded after data analysis was complete.

### Tandem affinity purification of xUB-conjugated proteins

8 dishes (10 cm in diameter) of GT1-7 cells were transfected with HBT-xUb/V5-xUbcH7and the xUba1-xUbch7-xRNF216 combination or xUba1-xUbch7-RNF216 CA cascade and collected after 24hr incubation. To inhibit proteasome activity, cells were treated with 10 μM MG132 for 4 h at 37°C. Cells were then washed twice with ice-cold 1 × PBS, pH 7.4, and harvested using a cell scraper with buffer A (8M urea, 300mM NaCl, 50mM Tris, 50mM NaH2PO4, 0.5% NP-40, 1mM PMSF and 125 U/ml Benzonase, pH 8.0). For Ni-NTA purification, cell lysates were centrifuged at 15,000 × g for 30 min at room temperature. 35 μL of Ni^2+^ Sepharose beads (GE Healthcare) for each 1mg of protein lysate was added to the clarified supernatant. After incubation OT at RT in buffer A with 10mM imidazole on a rocking platform, Ni^2+^ Sepharose beads were collected by centrifugation at 100 × g for 3 min and washed sequentially with 20-bead volume of buffer A (pH 8.0), buffer A-1 (pH 6.3), and buffer A-2 (pH 6.3) with 10mM imidazole. After washing the beads, proteins were eluted twice with 5-bead volumes of buffer B (8M Urea, 200mM NaCl, 50mM Na2HPO4, 2% SDS, 10mM EDTA, 100mM Tris, 250mM imidazole, pH 4.3). For streptavidin purification, the pH of the eluted fractions was adjusted to 8.0. A total of 50 μl of streptavidin-sepharose beads (Thermo Scientific, Rockford, IL) was added to the elution to bind ubiquitinated proteins. Samples were incubated on a rocking platform overnight at room temperature, streptavidin beads were collected and washed sequentially with 1.5 mL buffer C (8M Urea, 200mM NaCl, 2% SDS, 100mM Tris, pH 8.0), buffer D (8M Urea, 1.2MNaCl, 0.2% SDS, 100mM Tris, 10% EtOH, 10% Isopropanol, pH 8.0) and buffer E (8M urea, 100 NH4HCO3, pH 8).

### Sample digestion

Residual buffer E was removed and 200 μL of 50 mM NH4HCO3 was added to each sample, which were then reduced with dithiothreitol (final concentration 1 mM) for 30 min at 25 °C. This was followed by 30 min of alkylation with 5 mM iodoacetamide in the dark. The samples were then digested by adding 1 μg of lysyl endopeptidase (Wako) at RT for 2 h and further digested overnight with 1:50 (w/w) trypsin (Promega) at room temperature. Digested peptides were acidified with 25 μL of 10% (v/v) formic acid (FA) and 1% (v/v) triflouroacetic acid (TFA) and desalted with a Sep-Pak C18 column (Waters). Briefly, the Sep-Pak column was washed with 1mL of methanol and 1mL of 50% (v/v) acetonitrile (ACN) followed by equilibration which was performed with 2 rounds of 1mL of 0.1% (v/v) TFA in water. The acidified peptides were then loaded on a column washed with 2 rounds of 1 mL 0.1% (v/v) TFA. Elution was carried out by 2 rounds of 50% (v/v) ACN (400 μL each) and the resulting peptide eluent dried under vacuum.

### LC-MS/MS analysis

The data acquisition by LC-MS/MS was adapted from a published procedure^68^. Derived peptides were resuspended in the loading buffer (0.1% trifluoroacetic acid, TFA) and were separated on a Water’s Charged Surface Hybrid (CSH) column (150 µm internal diameter (ID) x 15 cm; particle size: 1.7 µm). The samples were run on an EVOSEP liquid chromatography system using the 15 samples per day preset gradient (88 min) and were monitored on a Q-Exactive Plus Hybrid Quadrupole-Orbitrap Mass Spectrometer (ThermoFisher Scientific). The mass spectrometer cycle was programmed to collect one full MS scan followed by 20 data dependent MS/MS scans. The MS scans (400-1600 m/z range, 3 x 10^6^ AGC target, 100 ms maximum ion time) were collected at a resolution of 70,000 at m/z 200 in profile mode. The HCD MS/MS spectra (1.6 m/z isolation width, 28% collision energy, 1 x 10^5^ AGC target, 100 ms maximum ion time) were acquired at a resolution of 17,500 at m/z 200. Dynamic exclusion was set to exclude previously sequenced precursor ions for 30 seconds. Precursor ions with +1, and +7, +8 or higher charge states were excluded from sequencing.

### Proteome Discoverer

Mass spectrometry data was analyzed according to a published protocol^69^. Spectra were searched using Proteome Discoverer 2.1 against the 2020 mouse UniProtKB/Swiss-Prot database (17,041 target sequences). Searching parameters included fully tryptic restriction, precursor mass tolerance (± 20 ppm), and fragment mass tolerance (± 0.05 Da). Serine, threonine, and tyrosine phosphorylation (+79.9663 Da), methionine oxidation (+15.99492 Da), asparagine and glutamine deamidation (+0.98402 Da) and protein N-terminal acetylation (+42.03670) were variable modifications (up to 3 allowed per peptide); cysteine was assigned a fixed carbamidomethyl modification (+57.021465 Da). Percolator was used to filter the peptide spectrum matches to a false discovery rate of 1%.

### Experimental design and statistical analysis

All experiments follow a between-subjects design. Statistical analysis was conducted using GraphPad prism as described in the text for each experiment. Non-parametric tests are used when the criteria for using parametric tests are not met. Data is represented as mean ± SEM with statistical significance set at 95%.

### Data Availability

All data and materials are available upon request from the lead contact author.

## Notes

### Competing Interest Statement

The authors have declared no competing interest.

### Summary of Updates

Figure 3 revised; Figure 4 revised; Figure 7 revised; Figure S5 revised; text updated for figure callouts in results and figure legends

