## SUPPLEMENTAL MATERIAL for "Substrate Profiling of RNF216 Uncovers a Translation-Linked OTUD4 Regulatory Axis"

#### FIGURE S1

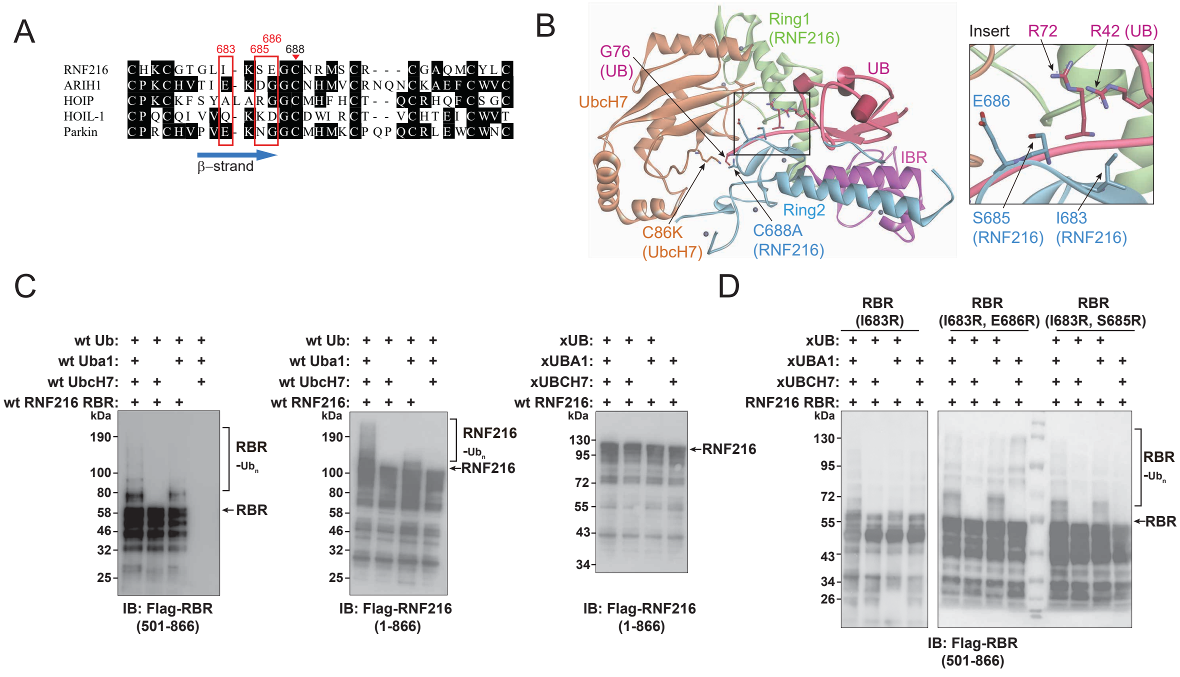


**Figure S1. Design and Validation of OUT for RNF216.**

(A) Sequence alignment of RBR E3s in the Ring2 region with residues in the β-strand of Ring2 highlighted in red frames. Combinations of I683R, S685R, and E686R mutations were incorporated into the RBR domain of RNF216 to assay if the mutations in the RBR domain could complement the R42E and R72E mutations in xUB to restore xUB transfer through the RBR.

(B) Crystal structure of the RBR domain of RNF216 bound to the UbcH7-Ub conjugate (PDB ID: 8EB0). The catalytic Cys residues in UbcH7 and the RBR domains were mutated (C86K in

UbcH7 and C688A in the RBR) to remove their reaction with Ub. The inset picture shows that residues anchored on the β-strand of the Ring2 domain of RNF216 RBR, including I683, S685, and E686, are in close vicinity to R42 and R72 in Ub that were mutated to Glu residues in xUB for its exclusive transfer through the OUT cascade.

(C) *Left two panels*, Wildtype (WT) Ub can be transferred through the WT RBR domain of RNF216 (residues 501-866) or full-length RNF216 (1-866) in the self-ubiquitination reaction. *Right,* WT RNF216 was not reactive with xUB transfer through the xUBA1-xUBCH7 pair to generate self-ubiquitinated RNF216.

(D) RBR mutants of RNF216 with the combinations of I683R, S685R, and E686R mutations could restore xUB transfer through the RBR mutants in the self-ubiquitination reaction.

**FIGURE S2**

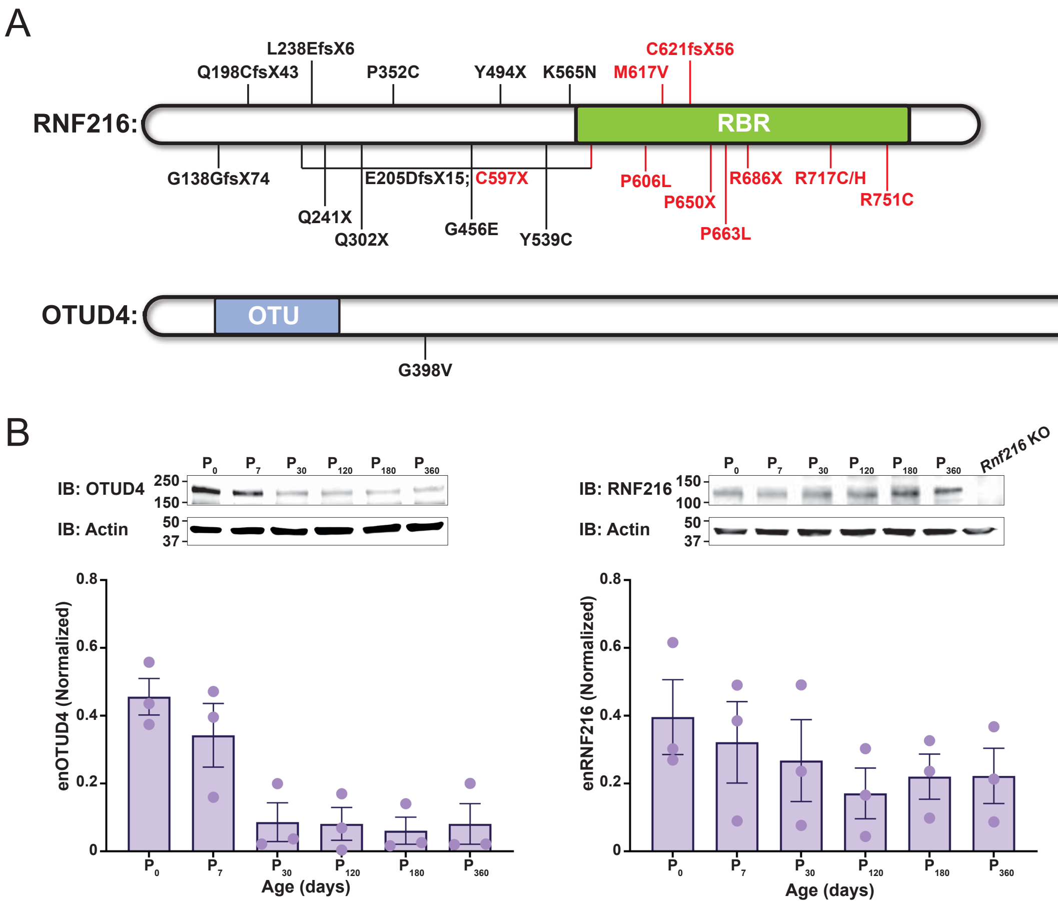
A
RNF216:
RBR
L238EfsX6
Q198CfsX43
P352C
Y494X
K565N
C621fsX56
M617V
G138GfsX74
E205DfsX15; C597X
Q241X
Q302X
G456E
Y539C
P606L
P650X
P663L
R686X
R717C/H
R751C
OTUD4:
OTU
G398V
B
IB: OTUD4
IB: Actin
IB: RNF216
Rnf216 KO
enOTUD4 (Normalized)
enRNF216 (Normalized)
Age (days)

**Figure S2: Gordon Holmes Syndrome is caused by mutations within the Ubiquitin machinery.**

(A) Mutations in the RBR E3 RNF216 and U-Box CHIP/Stub1 are found in GHS. A single point mutation in OTUD4 was identified in GHS.

(B) OTUD4 expression is decreased across development while RNF216 levels remain constant. Whole brain extracts from p0, p7, p30, p120, p180 and p360 were prepared from WT mice. Prepared samples were immunoblotted with anti-OTUD4 (left) or anti-RNF216 (right) antibodies with Actin used as a loading control. *Bottom*, Quantification of OTUD4 and RNF216 ** $p < 0.05$, *** $p < 0.005$. One-way ANOVA with Dunnett’s multiple comparisons test. N = 3 independent biological replicates.

### FIGURE S3

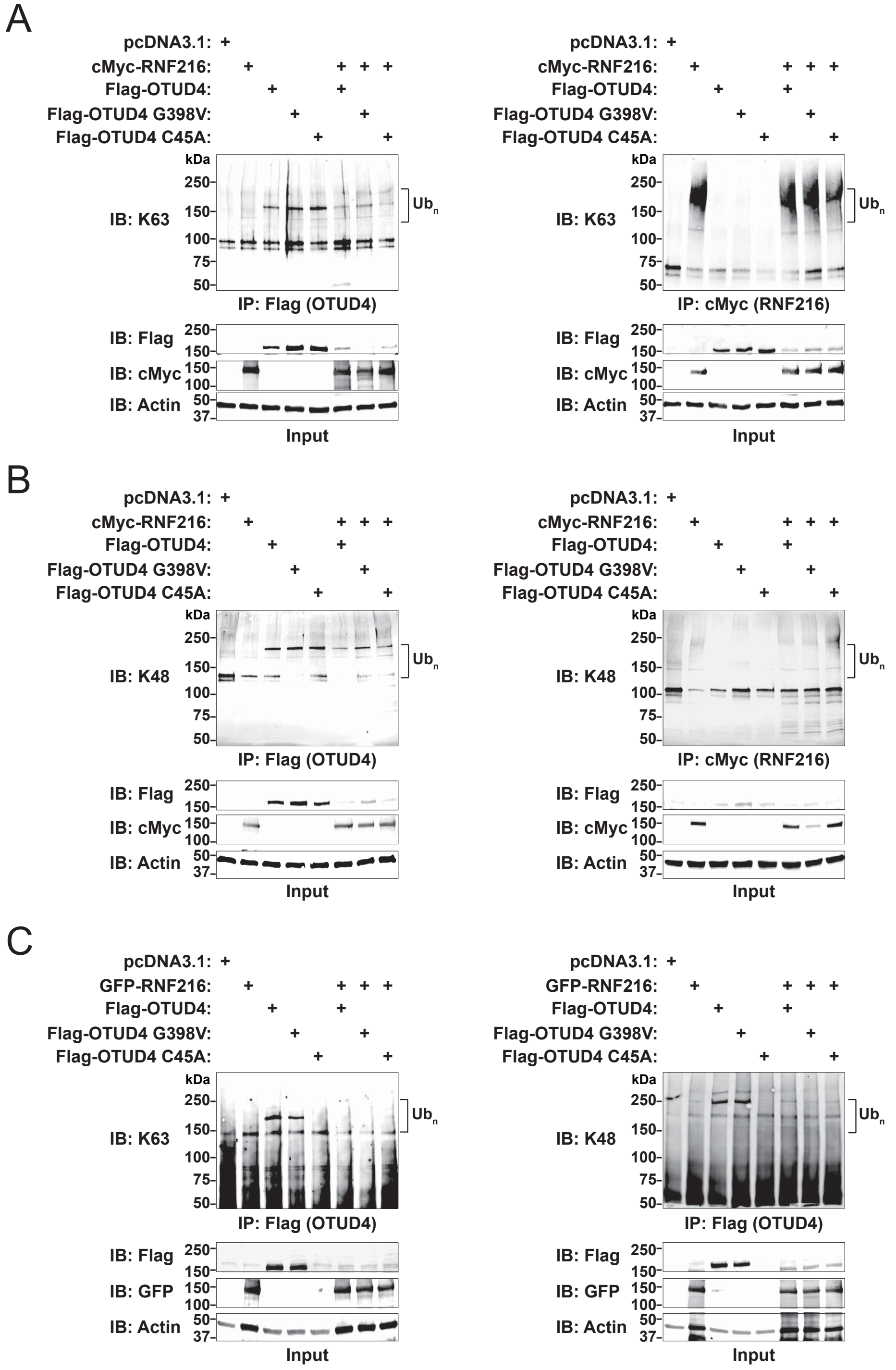
A
pcDNA3.1:
cMyc-RNF216:
Flag-OTUD4:
Flag-OTUD4 G398V:
Flag-OTUD4 C45A:
kDa
250-
150-
100-
75-
50-
IB: K63
Ub$_n$
IP: Flag (OTUD4)
IP: cMyc (RNF216)
IB: Flag
IB: cMyc
IB: Actin
37-
Input
B
IB: K48
C
GFP-RNF216:
IB: GFP

**Figure S3: RNF216 does not assemble homotypic K63- and K48- chain types on OTUD4.**

(A) RNF216-mediated ubiquitination of OTUD4 is not via K63-linkages. K63-specific Tandem Ubiquitin Binding Entities (TUBEs) were used to measure changes in OTUD4 ubiquitination (*Left*) and RNF216 autoubiquitination (*Right*).

(B) RNF216-mediated ubiquitination of OTUD4 is not via K48-linkages. K48-specific TUBEs were used to measure changes in OTUD4 ubiquitination (*Left*) and RNF216 autoubiquitination (*Right*).

(C) RNF216-mediated ubiquitination of OTUD4 is not K63- or K48-linkage specific as measured using Ub chain-specific antibodies for K63 (*Left*) or K48 (*Right*).

**FIGURE S4**

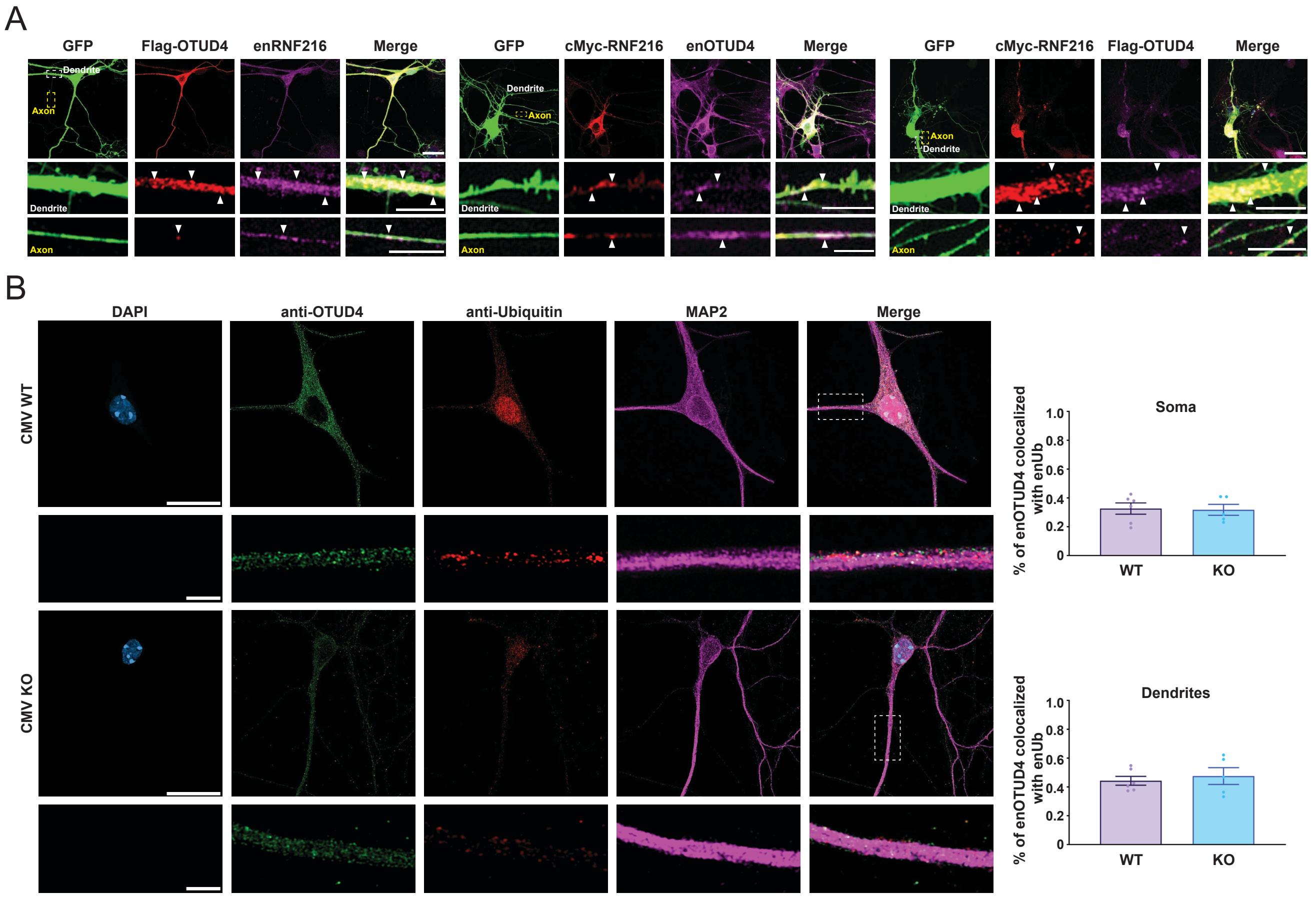


**Figure S4: RNF216 and OTUD4 co-localize in soma, dendrites and axons.**

(A) RNF216 and OTUD4 were found to colocalize in neuronal regions that include dendrites, axons, and soma. Primary hippocampal neurons were transfected at DIV13 with plasmids expressing GFP, myc-RNF216, and FLAG-OTUD4 in various combinations. Neurons were fixed 48 hours later and stained with antibodies for RNF216, OTUD4, Myc, or Flag. Myc-RNF216 was shown to colocalize with endogenous (en) OTUD4 and Flag-OTUD4 was shown to colocalize with endogenous (en) RNF216, particularly in dendrites.

(B) OTUD4 colocalization with Ubiquitin is unaffected in *Rnf216* KO cells. Primary hippocampal neurons from *Rnf216* WT or KO mice were fixed at DIV15 and stained with antibodies for OTUD4, UB, and Map2. *Left*, A fraction of Ubiquitin was shown to colocalize with endogenous (en) OTUD4. *Right*, Overlap quantification for Ub signal and (en) OTUD4.

#### FIGURE S5

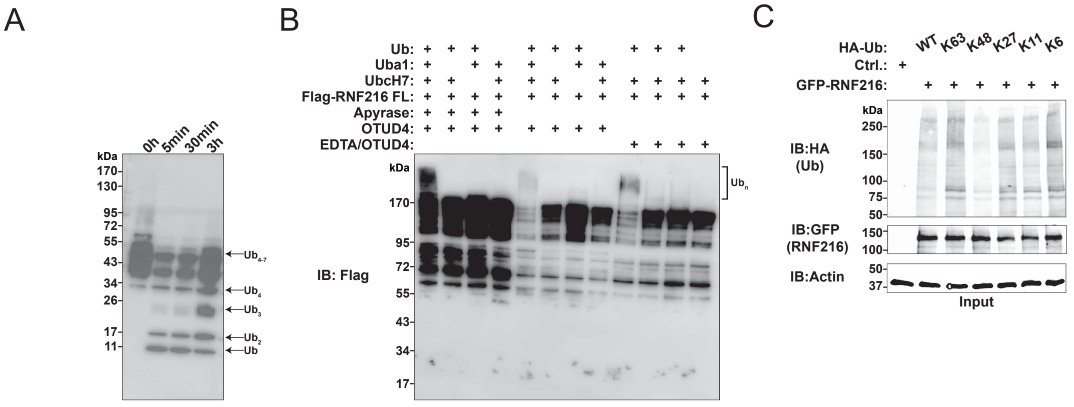


**Figure S5: OTUD4 deubiquitinates RNF216.**

(A) Purified OTUD4 (1-180) is catalytically active towards polymeric assembled Ub chains. Purified OTUD4 was subjected to polymeric Ub assembled chains for 5 min, 30 min, and 3h. Products were probed with an anti-UB antibody.

(B) OTUD4 (1-180) cleaves RNF216 ubiquitinated chains. Recombinant Flag-RNF216 was in vitro ubiquitinated. Following ubiquitination, Ub reactions were terminated with either Apyrase or EDTA and OTUD4 was then added to the reaction. In-vitro reaction products were probed with an anti-Flag antibody.

(C) Input blots from Figure 4E were immunoblotted with anti-GFP or -Actin antibodies.

**FIGURE S6**

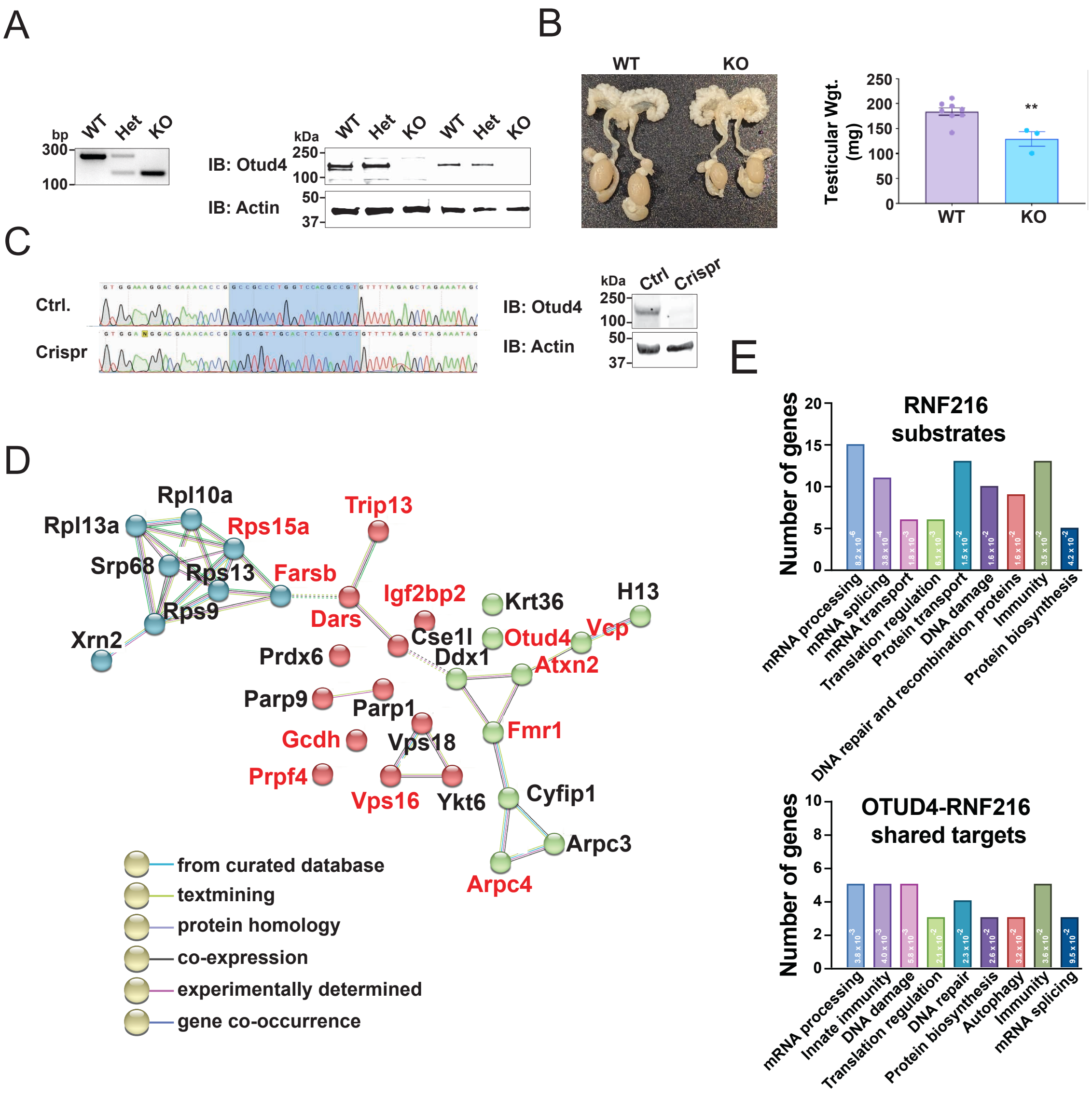
A
bp
WT
Het
KO
300
100
kDa
250
100
50
37
IB: Otud4
IB: Actin
B
WT
KO
Testicular Wgt. (mg)
250
200
150
100
50
0
**
C
Ctrl.
Crispr
IB: Otud4
IB: Actin
Ctrl
Crispr
D
Rpl13a
Rpl10a
Rps15a
Trip13
Srp68
Rps13
Farsb
Igf2bp2
Rps9
Dars
Krt36
H13
Xrn2
Cse1l
Otud4
Vcp
Prdx6
Ddx1
Atxn2
Parp9
Parp1
Gcdh
Vps18
Fmr1
Prpf4
Vps16
Ykt6
Cyfip1
Arpc3
Arpc4
from curated database
textmining
protein homology
co-expression
experimentally determined
gene co-occurrence
E
RNF216 substrates
Number of genes
20
15
10
5
0
mRNA processing
mRNA splicing
mRNA transport
Translation regulation
Protein transport
DNA damage
DNA repair and recombination proteins
Immunity
Protein biosynthesis
OTUD4-RNF216 shared targets
10
8
6
4
2
0
mRNA processing
Innate immunity
DNA damage
Translation regulation
DNA repair
Protein biosynthesis
Autophagy
Immunity
mRNA splicing

**Figure S6: OTUD4 Crispr cell and KO mouse validation.**

(A) *Left*, Genotyping results from *Otud4*$^{+/+}$ (WT), *Otud4*$^{-/+}$ (HET) and *Otud4*$^{-/-}$ (KO) mice. *Right*, Representative Western blots for OTUD4 in male (left) and female (right) *Otud4*$^{+/+}$, *Otud4*$^{-/+}$, and *Otud4*$^{-/-}$ mice whole brain tissue lysates.

(B) Representative image of male reproductive organs in 90d old *Otud4* WT and KO mice. KO mice showed a significant reduction in testicular weights compared WT. Unpaired t-test ** p = 0.0406, N = 8 WT and 3 KO animals per group.

(C) *Top*, Sequence alignment in Control (Ctrl) and *Otud4* Crispr demonstrates sequence mismatch within the *Otud4* gene. *Bottom*, Representative Western blot for OTUD4 in GT1-7 CRISPR-Cas9 control and *Otud4* knockout (KO) cells.

(D) RNF216 substrate overlap with interaction partners of the OTUD4 interaction network identified by conducting a meta-analysis of OTUD4 binding partners from the Bio-Grid and previous published works. Proteins colored in red are known to be mutated in a neurological disease.

(E) *Top*, DAVID analysis predicting most significant function of 174 RNF216 substrates for GT1-7 RNF216 OUT screen. *Bottom*, DAVID analysis of the 36 RNF216-OTUD4 common targets.

#### FIGURE S7

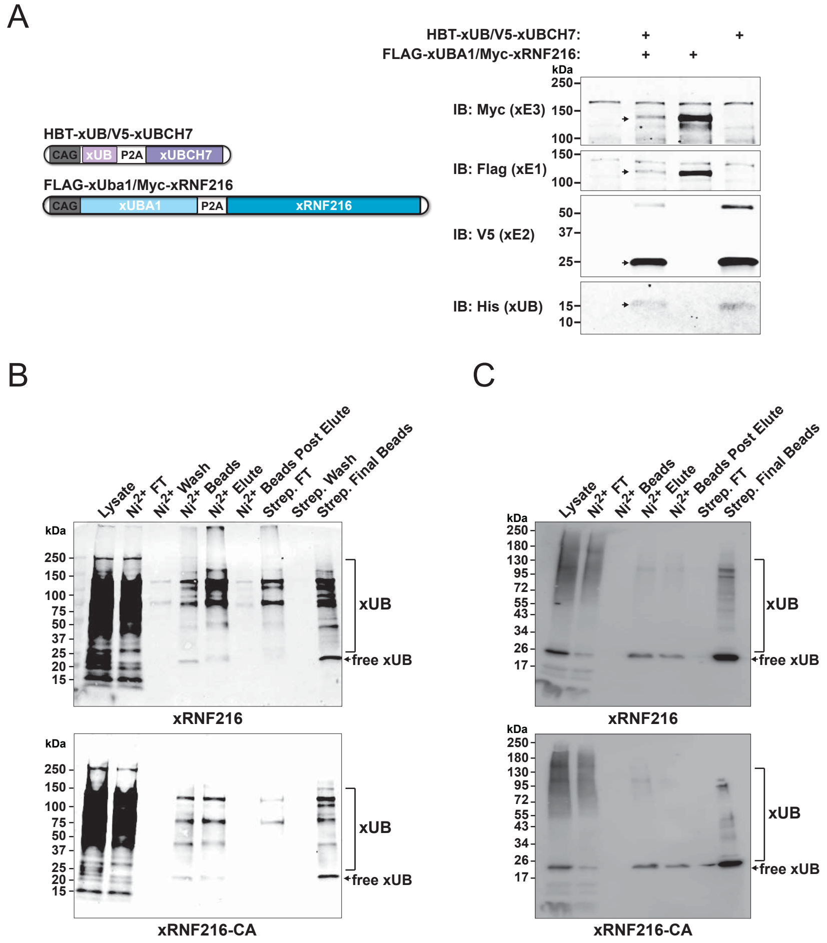


**Figure S7: RNF216 OUT identifies distinct substrates in HEK293 cells.**

(A) *Left*, Schematic of dual expression 2-plasmid system for expression of RNF216 OUT cascades in living cells. *Right,* Expression of OUT components and HBT-xUB upon co-transfection in the immortalized hypothalamic GT1-7 cell line.

(B) Validation of xUB conjugated purified proteins through sequential affinity columns of Ni-NTA and Streptavidin from the lysate of GT1-7 cells expressing the xRNF216 OUT cascade, N = 5 independent biological replicates. Purified targets were digested by trypsin and identified by LC-MS/MS.

(C) xUB conjugated proteins were purified from OUT cells expressing the xUBA1-xUBCH7-xRNF216 cascade enzymes with HBT-xUB (*top*) and from control cells expressing xUBA1-xUBCH7 pairing with xRNF216-C688A mutant and HBT-xUB (*bottom*).

**FIGURE S8**

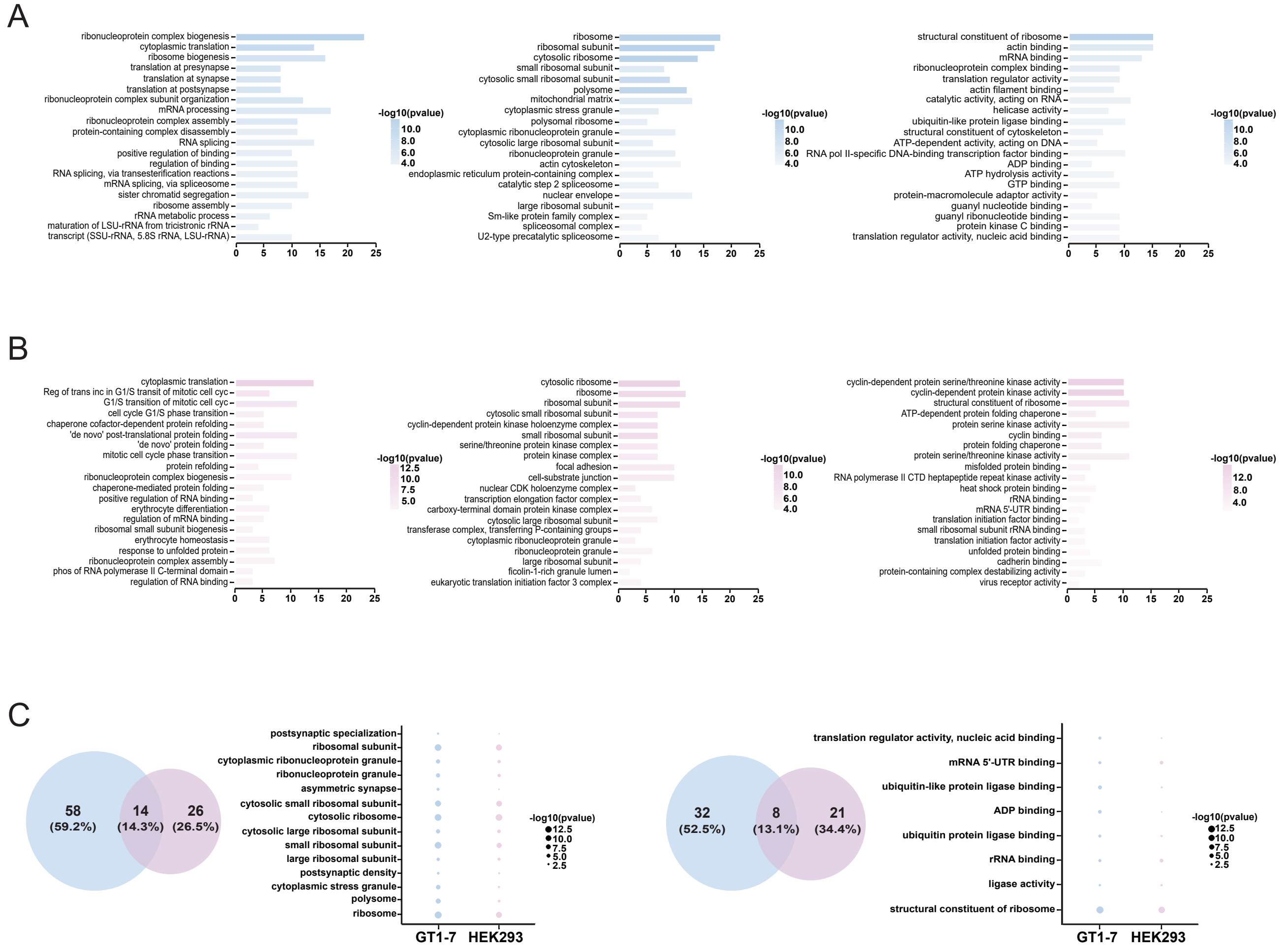


**Figure S8: Functional enrichment analysis for RNF216 OUT substrates from GT1-7 and HEK293 cells.**

(A) Top significant terms from list in GT1-7 cells of gene ontology related to Biological Process (*Left*), Cellular Component (*Middle*) and Molecular Function (*Right*).

(B) Top significant terms from list in HEK293 cells of gene ontology related to Biological Process (*Left*), Cellular Component (*Middle*) and Molecular Function (*Right*).

(C) Functional annotation of most significant shared terms related Cellular Component (*Left*) and Molecular Function (*Right*) of RNF216 substrates between HEK 293 and GT1-7 cells.

FIGURE S9

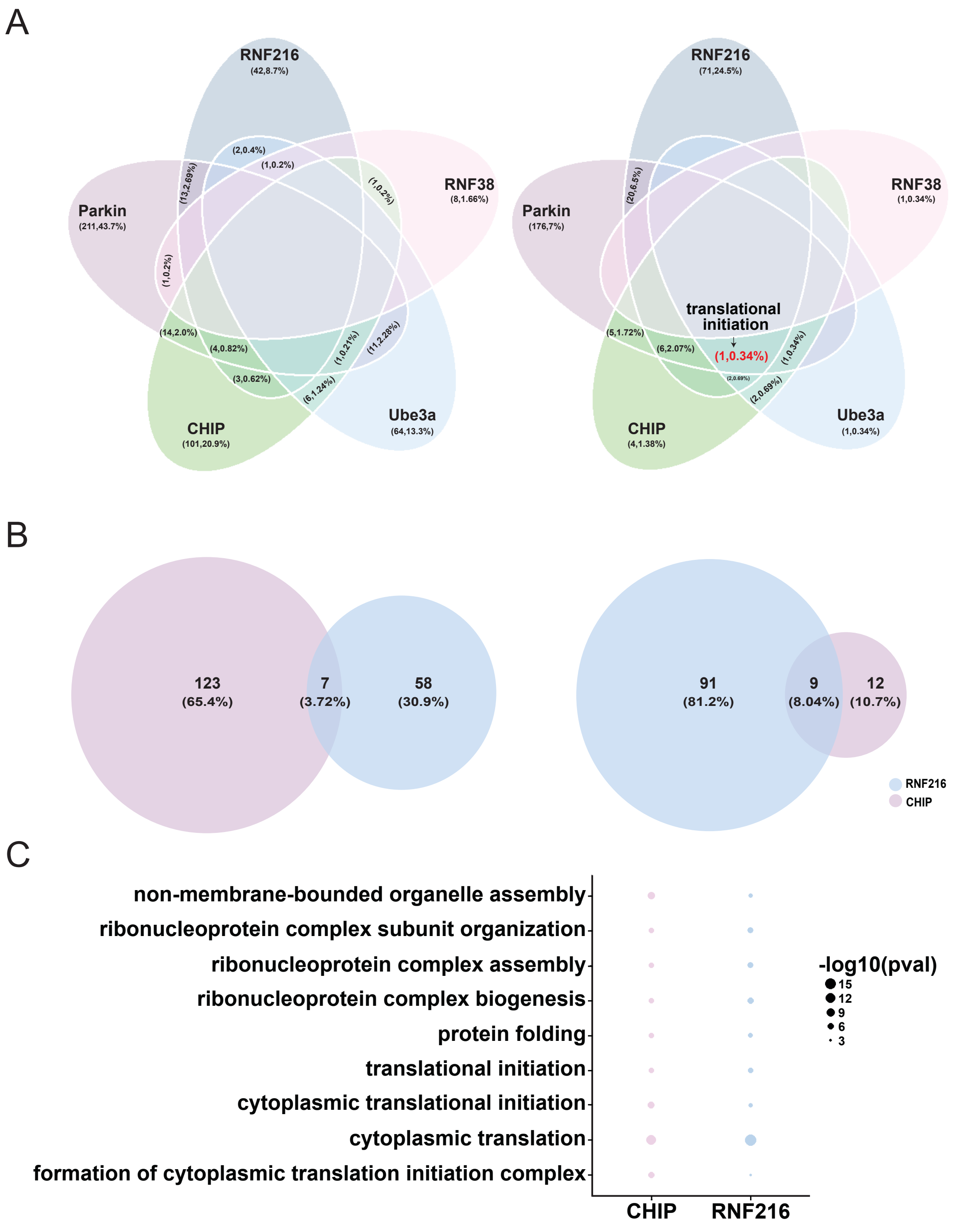
A
RNF216
(42,8.7%)
RNF38
(8,1.66%)
Parkin
(211,43.7%)
CHIP
(101,20.9%)
Ube3a
(64,13.3%)
RNF216
(71,24.5%)
RNF38
(1,0.34%)
Parkin
(176,7%)
CHIP
(4,1.38%)
Ube3a
(1,0.34%)
translational initiation
(1,0.34%)
B
123
(65.4%)
7
(3.72%)
58
(30.9%)
91
(81.2%)
9
(8.04%)
12
(10.7%)
RNF216
CHIP
C
non-membrane-bounded organelle assembly
ribonucleoprotein complex subunit organization
ribonucleoprotein complex assembly
ribonucleoprotein complex biogenesis
protein folding
translational initiation
cytoplasmic translational initiation
cytoplasmic translation
formation of cytoplasmic translation initiation complex
CHIP
RNF216
-log10(pval)
15
12
9
6
3

**Figure S9: Classes of E3s may regulate a suite of substrates involved in protein synthesis regulation subserving roles as proteostats.**

(A) *Left*, Comparison of substrates for RNF216, Parkin, RNF38, CHIP and UBE3A in HEK293 cells. Substrate overlap of 5 E3s from HEK 293 cells reveals common and uncommon substrates. *Right*, Functional annotation of shared biological function of substrate pools from E3 ligases. 4/5 E3 ligases share only 1 common biological function, which is translation initiation (GO:0006413).

(B) Comparison of substrates of RNF216 and CHIP in HEK cells. *Left*, Substrate overlap of RNF216 and CHIP substrates from HEK 293 cells reveals only 7 common substrates. *Right*, Functional annotation of shared biological functions of RNF216 and CHIP substrates.

(C) Functional annotation of most significant shared biological functions of CHIP and RNF216 substrates reveals protein translation as a common factor.
